# Spatiotemporal isoform profiling identifies a central activator of microexon splicing in *C. elegans*

**DOI:** 10.64898/2026.09.14.751627

**Authors:** Bina Koterniak, Ernest Liang, Michael Zoberman, Congrong He, Bikash Choudhary, Nour H. Sadek, Pallavi P. Pilaka-Akella, Adam D. Norris, John A. Calarco

## Abstract

Dynamic alternative splicing programs drive tissue specification and organismal development. Here, we present the most comprehensive, tissue-resolved alternative splicing dataset in *Caenorhabditis elegans* to date, spanning three major tissue types from early embryogenesis to adulthood. We uncovered broad developmental and tissue-regulated alternative splicing trends and identified putative RNA-binding proteins (RBPs) coordinating co-regulated splicing networks governing tissue identity. Among the network of regulated splice variants, we uncovered additional, unannotated microexons for further study in *C. elegans*. Through a forward genetic screen, we identified RBM-25, a U1 snRNP-associated regulatory protein, as a key regulator that preferentially enhances the inclusion of short exons and microexons alongside its binding partner PRP-40. Phenotypic profiling of *rbm-25* mutants demonstrated significant defects in behviour, reproductive fitness, lifespan, and synaptic signalling, indicating a key role for RBM-25-regulated target transcripts in normal physiology and behaviour. Together, these findings expand our appreciation for spatiotemporal isoform diversity during animal development, and provide mechanistic insights into how microexon splicing is regulated.

## Introduction

Alternative splicing (AS), a gene regulatory layer generating multiple mature messenger RNA (mRNA) isoforms from a single gene locus, is highly prevalent in metazoans and contributes to the differentiation of cells into various specialized tissues (Marasco & Kornblihtt, 2023; Ule & Blencowe, 2019; Wright et al., 2022). Levels of AS in eukaryotic genomes correlate with organismal cellular diversity (Chen et al., 2014), with humans having alternative isoforms in approximately 95% of multi-exonic genes (Pan et al., 2008; E. T. Wang et al., 2008), while in *C. elegans* that number is closer to 25% (Ramani et al., 2011). The process of mRNA splicing involves the stepwise assembly of a large (>3 MDa) ribonucleoprotein complex, the spliceosome, across a pre-mRNA transcript resulting in the catalytic removal of introns and the joining of exons (Wilkinson et al., 2020).

The core components of the spliceosome are the U1, U2, and U4/5/6 snRNPs, all characterized by strong evolutionary conservation, yet none of them harbours a pre-existing catalytic site for intron excision (Ohi, 2017). As a result, spliceosome assembly is a highly dynamic process that can be regulated by numerous splicing factors, which can jointly affect the activity and efficiency of the splicing at particular splice sites (Wahl et al., 2009a). Splicing factors, also referred to as *trans*-acting factors, are in turn recruited through sequence features known as *cis*-acting elements, which can be found in exonic or intronic regions and act as splicing enhancers or silencers, provide additional layers of regulatory control (Marasco & Kornblihtt, 2023). While alterations to *cis*-acting elements can lead to local splicing modifications, perturbations of splicing factors, including RNA-binding proteins (RBPs), can trigger significant perturbations across networks of co-regulated splicing events (Wong et al., 2017). Importantly, it has been estimated that close to 25% of human diseases are attributed to splicing misregulation (Padgett, 2012; Ward & Cooper, 2010), and this estimate may be conservative given ongoing discoveries of mutations impacting *cis*-elements or splicing factors (Dvinge, 2018). Given its importance, continued research into splicing regulatory mechanisms is required.

Alternative splice isoforms can exhibit variation in both their transcript and protein-level characteristics. Transcript export, subcellular localization, stability, and translational efficiency can be regulated through the AS-dependent inclusion of the appropriate signals or the coupling of AS to other downstream processes such as nonsense-mediated decay (NMD) and microRNA (miRNA) targeting (Hackl et al., 2023; Lewis et al., 2003; Müller-McNicoll et al., 2016; Zeng & Hamada, 2020). Alternative splicing can also modulate protein function by adjusting domain architecture, enzymatic/electrophysiological properties, and interaction networks of proteins (Kelemen et al., 2013; Kjer-Hansen & Weatheritt, 2023). Protein isoforms that are subject to tissue- and developmental regulation can display significantly distinct patterns in expression, functionality, and interaction profiles, thereby playing a pivotal role in shaping tissue and developmental states (Baralle & Giudice, 2017; Kalsotra & Cooper, 2011; Yang et al., 2016). Given the important roles for tissue- and developmentally-biased isoforms, it remains an important endeavour to map these splice variants in multicellular animals.

An interesting class of spatiotemporally-regulated alternative splicing events involve microexons, very short exons that can measure as few as 3nt and yet often have important roles in protein function (Gonatopoulos-Pournatzis & Blencowe, 2020; Mackensen & Irimia, 2025; Scheckel & Darnell, 2015). While early studies highlighted the function and splicing dynamics of individual microexons (Black, 1991; Inman et al., 1998; McAllister et al., 1992; Santoni et al., 1989; Small et al., 1988), recent advances in RNA-sequencing technologies and downstream analytical pipelines have uncovered widespread microexon splicing (Irimia et al., 2014a; Y. I. Li et al., 2015a; Ustianenko et al., 2017). Subsequent research has established the pivotal role of microexons in the tissue-specific functions of predominantly the nervous system, including neurons and microglia (Irimia et al., 2014b; Lee et al., 2020; Y. I. Li et al., 2015b), but also in the muscle, heart, retina and pancreas (Ciampi et al., 2022; Juan-Mateu et al., 2023; Murphy et al., 2016; Parada et al., 2020; W. Wang et al., 2024).

Microexons exhibit a strong tendency toward reading frame preservation, for example approximately 80% of microexons in humans are multiples of 3nt (Irimia et al., 2014b; Y. I. Li et al., 2015b). Furthermore, microexons and flanking introns are highly evolutionarily conserved, with microexons frequently embedded within or adjacent to conserved protein domains, disordered regions or surface residues, where they modulate protein functions and interactions (Irimia et al., 2014b; Y. I. Li et al., 2015b; Roth et al., 2023; Torres-Méndez et al., 2022).

Mechanistically, due to their small size, the standard “exon definition” model for splicing that is favoured when large introns flank small exons does not readily support the inclusion of microexons (Sterner et al., 1996). Allosteric hindrance prevents the large spliceosome from assembling at the 5’- and 3’-splice sites, and this hindrance can be relieved when the microexon has been artificially expanded with no additional changes to the flanking introns (Black, 1991). Microexons are also too small to accommodate many exonic-splicing enhancers (ESEs) (Curry-Hyde et al., 2018; Ustianenko et al., 2017). However, multiple compensatory properties of microexons and flanking sequences work to offset these deficits. For example, in the human brain it has been demonstrated that constitutive microexons have a higher density of ESEs and ISEs (intronic splicing enhancers) and stronger 5’- and 3’-splice sites (Carlo et al., 2000; Y. I. Li et al., 2015b). AS microexons have an enrichment of pyrimidines at 10-20nt upstream of the 3’-splice site, possibly bound by U2AF and subsequently U2 snRNP, or several other RBPs that have U/C rich binding motifs (Y. I. Li et al., 2015b; Ustianenko et al., 2017).

In vertebrates, neuronal microexon inclusion is largely regulated by SRRM4/nSR100 and its paralog, SRRM3 (Calarco et al., 2009; Gonatopoulos-Pournatzis et al., 2018). Mechanistically, SRRM3/4 bind to intronic UGC-motifs close to the microexon 3’-splice site to promote early spliceosome recruitment. A specialized domain called eMIC (enhancer of microexons), found in the C-term of SRRM3/4, is necessary and sufficient to drive microexon inclusion (Torres-Méndez et al., 2019). Additionally, the splicing cofactors SRSF11 and RNPS1 interact with SRRM3/4 at intronic splicing enhancer elements to further stabilize early spliceosome assembly, particularly the U2 snRNP, thereby facilitating accurate exon definition and efficient splicing of neuronal microexons (Gonatopoulos-Pournatzis et al., 2018). However, we still lack a complete picture of how microexons are included in specific tissue- and developmental-specific contexts.

Increasingly rich *C. elegans* transcriptome datasets reveal a complex landscape of alternative splicing regulation across developmental stages and tissues, enabled by a technological evolution from microarrays (Barberan-Soler et al., 2009; Barberan-Soler & Zahler, 2008) to high-throughput RNA-seq (Gerstein et al., 2010; Ramani et al., 2011), and from whole-animal (Tourasse et al., 2017) to tissue- and cell-specific resolution (Kaletsky et al., 2018; Koterniak et al., 2020; Warner et al., 2019). Parallel to splicing patterns seen in vertebrates (Barbosa-Morais et al., 2012; Yeo et al., 2004), the worm nervous system demonstrates the highest levels of alternative splicing relative to other tissues (Koterniak et al., 2020). Recent studies of alternative splicing in the *C. elegans* nervous system, enabled by deep, cell-specific transcriptomes from the CeNGEN consortium, have also revealed unique splicing patterns across neuronal cell types (Weinreb et al., 2025; Wolfe et al., 2025).

Microexons have also been previously detected in *C. elegans* (Koterniak et al., 2020; Torres-Méndez et al., 2019; Volfovsky et al., 2003; Weinreb et al., 2025). However, a systematic genome-wide search using targeted computational pipelines to comprehensively identify *C. elegans* microexons has not been conducted. Additionally, the molecular mechanisms governing microexon regulation in worms are only beginning to be elucidated. PRP-40 has been demonstrated to be a major activator of microexon inclusion in worms (Choudhary et al., 2021), and its ortholog, PRPF40A, appears to co-regulate microexon inclusion with SRRM4 in mouse neuroblastoma cells (Choudhary & Norris, 2025). Nonetheless, an unbiased search for regulators of microexon splicing has yet to be performed. This gap in knowledge is underscored by the absence of an eMIC domain in worm SR-related protein homologs, which raises questions regarding the alternative mechanisms that activate microexon inclusion, particularly within the nervous system.

In this study, we performed the most extensive meta-analysis of *C. elegans* spatiotemporal alternative splicing patterns to date, leveraging a comprehensive RNA-seq dataset encompassing tissue- and developmental stage-specific information for three major tissue types during embryogenesis, late larval development, and adulthood. Our findings illuminate the complex interplay between various regulatory mechanisms governing alternative splicing outcomes in tissues and across development, and highlight several key splicing programs for further study. We further used a specialized discovery pipeline to perform a genome-wide search for both annotated and novel microexons, effectively doubling their count within *the C. elegans* genome. Guided by this important class of splicing events, we performed a genetic screen to identify novel regulators of neuronal microexon splicing, uncovering the conserved RNA-binding protein, RBM-25, as a central activator of these exons. Additionally, we demonstrate that RBM-25 and PRP-40 physically interact with each other, and loss of either factor leads to pronounced global effects on microexon splicing. Finally, we also demonstrate that loss of RBM-25 leads to brood size, lifespan, synaptic signalling, and behavioural defects, highlighting the physiological importance of the program of targets regulated by this factor. Taken together, our results highlight the importance of alternative splicing in shaping tissue identity during development, and provide mechanistic and phenotypic insight into an important program of regulated exons.

## Results

### Transcript abundance and alternative splicing both play key roles in defining the transcriptomes of three major C. elegans tissues throughout development

Expanding upon previous studies of steady-state transcript levels and alternative splicing in *C. elegans* across tissues at individual developmental stages, we consolidated a set of three deeply sequenced datasets from embryos (Warner et al., 2019), fourth larval stage (L4; (Koterniak et al. 2020) and adults (Kaletsky et al., 2018), profiling the transcriptomes or ribosome-associated mRNAs of three major tissues: intestine, body-wall muscle, and neurons (see Methods for details). We evaluated the gene expression and alternative splicing programs in each tissue over developmental time, hypothesizing that a more comprehensive meta-analysis would provide insight into the contribution of each aspect of gene regulation to tissue development. We first performed PCA analyses on standardized log2-transformed normalized gene counts as a measure of steady-state transcript levels (referred to hereafter as gene expression (GE)), and PSI (percent spliced in) values of local splice variations (LSVs) (**Fig. 1** and **Supplemental Fig. S1**; see Methods for details). The embryonic data were also analyzed separately from the L4/Adult data in order to study early developmental transitions across tissues at a higher resolution.

**Figure 1:**
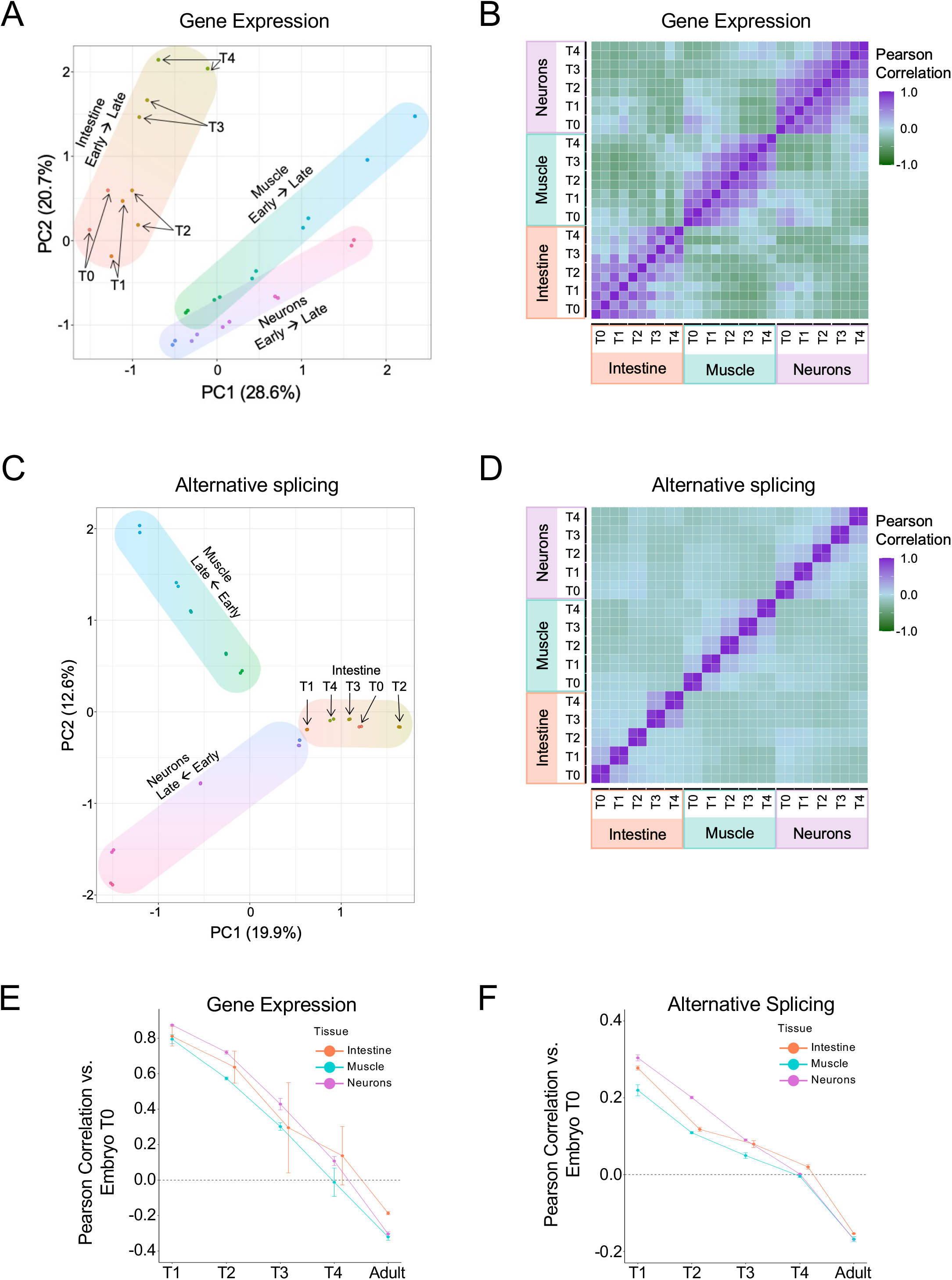
Tissue-specific gene expression and alternative splicing profiles across embryonic developmental time points. A) Principal Component Analysis (PCA) plot of normalized gene expression counts (log₂-transformed, standardized) at five embryonic time points T0 (early) to T4 (late), two replicates each, in three tissues: intestine (orange–green), muscle (green– blue), and neurons (purple–pink). Arrows indicate the direction of early to late time points. B) Matrix of pairwise Pearson correlations of the gene expression data described in A). Each cell shows the correlation between two samples, with purple indicating strong positive correlation and green, strong negative correlation. C) PCA plot of standardized percent spliced in (PSI) values of all the alternative splicing events detected in the data described in A) Arrows indicate the direction of early to late time points. D) Matrix of pairwise Pearson correlations of the alternative splicing data described in C). Each cell shows the correlation between two samples, with purple indicating strong positive correlation and green, strong negative correlation. E-F) Pearson correlation values comparing the early embryonic timepoint T0 with every subsequent time point to examine the trajectory of E) gene expression pattern and F) alternative splicing pattern over embryonic development. Lines show the mean Pearson correlation between matched replicates, and error bars represent ± SD across replicate correlations.

The *C. elegans* embryo contains 558 cells with invariant lineages (Sulston et al., 1983; Sulston & Horvitz, 1977). The embryo RNA-seq data by Warner et al., 2019 uses FACS (fluorescence-activated cell sorting) sorted cells to sequence five time points (T0-T4). The initial time point captures embryos halfway through gastrulation (∼150 min post first cleavage (pfc)) then at 90 min intervals (T1:250 min pfc; T2:340 min pfc; T3:430 min pfc) until the last time point, T4, as the embryo reaches the three-fold stage at 520 min pfc. In this timeframe, myoblasts (body-wall muscle precursor cells) arise upon the completion of gastrulation at ∼290 min (between T1 and T2) and then migrate to quadrants of the worm circumference. By ∼350 min (just prior to T2) all muscle cells have completed divisions (81 cells) and reached their final positions where they will polarize and change in morphology and physical linkage. Fourteen additional body-wall muscle cells arise post-embryonically. Similarly, 222 out of 302 neurons are generated during embryogenesis in a burst of cell divisions spanning from ∼280-410 minutes (between T1 and T3) and usually migrate short distances to their final location (Sulston et al., 1983). The remaining neurons are born in the L1 and L2 larval stages. Neurite outgrowth and the formation of synapses starts from the embryonic stages and continues into adulthood to form the connectome (Cook et al., 2019; Godini et al., 2022; Varier & Kaiser, 2011). The expression profiles of neurons derived from similar lineages may diverge as they acquire their terminal fates, whereas neurons originating from distinct lineages may exhibit developmental convergence as they adopt similar terminal identities (Packer et al., 2019; Poole et al., 2024). In contrast, the intestine is undergoing rapid changes very early in embryogenesis. By the time the T0 timepoint occurs at ∼150 min, endodermal precursor cells, Ea and Ep, have already internalized and undergone coordinated divisions and migrations (Bucher & Seydoux, 1994). Between T0 and T2 (∼120-300min), the developing intestine will expand from 4 to 16 cells and undergo a final set of divisions to bring the number of intestinal cells to 20 (Sulston & Horvitz, 1977).

PCA analysis of embryonic tissue-specific GE and AS patterns confirmed that samples had a tendency to cluster closely with their corresponding tissues, and the two replicates within each timepoint also cluster most closely to each other (**Fig. 1A** and **1C**). This indicates that the principal components measured in both gene expression (GE) and alternative splicing (AS) data effectively capture variation present in the high-dimensional dataset. In muscle and neurons, GE and AS patterns presented in the PCA plots also reflect developmental continuity, with samples ordering according to developmental timepoint from early (T0) to late (T4), suggesting that progressive and dynamic transcriptional programs are being established. Also, in most cases, there is a large separation between early and late embryo timepoints, indicating that development is a major source of transcriptional variation in embryos. Of the three tissues, the intestine exhibits the lowest degree of replicate clustering in the GE PCA (**Fig. 1A**) and a lack of temporal continuity across developmental timepoints in the AS PCA (**Fig. 1C**). The intestine GE data is also the most separate from the other two tissues in the PCA space, and we find that the intestine has the highest number of tissue-regulated, differentially expressed genes (DEGs) across all embryonic timepoints (intestine: 11231; muscle: 10491; neurons: 10525; log_2_ fold change of at least 1.5; **Supplemental Table S1**). Single-cell RNA-seq (sc-RNA-seq) of *C. elegans* embryos revealed that intestinal cells reach terminal differentiation relatively early compared to body-wall muscle and neurons, but intestinal cell type-specific transcriptomes remain highly similar even after specification (Packer et al., 2019). These results are consistent with our observations that the intestine has the fewest number of DEGs showing developmental regulation within the embryo stages profiled (intestine: 3952; muscle: 7011; neurons: 5892; **Supplemental Table S1**).

We also generated Pearson correlation matrices using the same GE and AS data to quantify the similarity between any two samples in the embryonic data. The GE correlation matrix highlights the robust correlations in GE patterns within each specific tissue, as seen by the three discrete blocks of high correlation (**Fig. 1B**). As expected, the correlation in GE decreases with increasing distance between developmental timepoints. In contrast, the alternative splicing (AS) correlations reveal a distinct pattern, where high correlations are predominantly observed among the two replicates within each timepoint (**Fig. 1D**). Outside of the replicate similarity, there is very little correlation between samples, even in other timepoints within the same tissue. This suggests that the splicing features underpinning each developmental timepoint demonstrate highly dynamic temporal regulation. These findings are consistent with studies in mammalian systems showing that alternative splicing patterns can provide resolution equal to or greater than gene expression profiles in distinguishing neuronal subtypes and cancer states (Feng et al., 2021; Ha et al., 2021; Stricker et al., 2017; Zhang et al., 2013).

We also compared GE and AS data in L4 and adult stages (**Supplemental Fig. S1**). The PCA plot of AS values (**Supplemental Fig. 1C**) created more clearly defined clusters for the three tissues compared to the GE data (**Supplemental Fig. 1A** and **1C**). Within each tissue, there is also some separation between the L4 and adult stage samples in the GE and AS plots. All three tissues are undergoing maturation to their final form and function, including changes associated with the onset of reproduction and sex-specific traits (Altun & Hall, 2009; Ghaddar et al., 2023). A recent study investigated transcriptional changes between L4 and adult stage neurons and identified gene expression differences that support the emergence of adult-specific functions related to learning/memory and behaviour (St. Ange et al., 2024).

Correlation analyses revealed that gene regulatory patterns differ substantially between L4 and adult stages across all tissues. This divergence is clear in the GE data (**Supplemental Fig. 2B**) and becomes even more pronounced in the AS data (**Supplemental Fig. 2D**). Although these trends likely reflect *bona fide* transcriptome differences in these late developmental periods, it remains unclear to what extent these stage-specific differences reflect real transcriptomic changes rather than technical variation caused by differing profiling approaches used to generate the L4 and adult datasets. For this reason, in this study, we decided to avoid direct comparisons between the L4 data and the embryo/adult data.

We also probed the divergence of GE and AS data over developmental time in each tissue by computing the correlation between each developmental stage (excluding L4) with the early embryonic timepoint (T0) (**Fig. 1E** and **1F**). In both cases there was a progressive divergence of each tissue from the early embryonic stage. While GE at later embryonic timepoints maintained a relatively high correlation to the early embryonic timepoint until the late embryo and adult stages, AS correlation to T0 was consistently lower. This suggests that GE changes occur gradually, with early embryonic expression patterns remaining active and highly correlated until later in development. Conversely, AS profiles diverge rapidly, showing a sharp early decline in correlation to T0 that highlights how splicing patterns are highly unique to each specific developmental stage.

Collectively, our meta-analysis provides the most comprehensive map to date of spatiotemporal isoform utilization across broad tissues in *C. elegans*. Our analysis further indicates that both gene expression (GE) and alternative splicing (AS) regulation play pivotal roles in tissue specification but have distinct dynamics over the course of development.

### Developmentally regulated splicing programs form coherent networks and shape tissue identity

Given the previous observation that embryonic splice isoforms create unique AS profiles in the three broad tissue types, we next sought to explore the relationship between developmentally-regulated (D-R) and tissue-regulated (T-R) AS (**Fig. 2** and **Supplemental Fig. S2**). D-R events refer to splicing events exhibiting significantly differential usage patterns between two developmental stages within the same tissue. For this D-R isoform analysis, we considered early embryo vs. late embryo and late embryo vs. adult stages as key transition points for comparison. Conversely, T-R events involve splicing events with differential usage patterns between two tissues at the same developmental stage, which also included measurements from L4 data. Our differential splicing analysis (see Methods for details) identified 1,702 unique D-R events and 2,834 unique T-R events (delta PSI; |dPSI| >= 0.20; **Supplemental Table S2)**. To compare the relationship between D-R and T-R splicing events, we visualized these data in a series of heatmaps and Venn Diagrams (**Fig. 2A**, **Supplemental Fig. 2A** and **2D**). Notably, for the majority of these comparisons, a prevailing trend emerged where the same LSVs displaying significant usage differences between developmental transitions within a given tissue were also tissue-defining (**Fig. 2A** and **2B**, neurons as reference tissue). Similar trends were observed for comparisons anchored by the other two tissues (**Supplemental Fig. 2A-D**).

**Figure 2:**
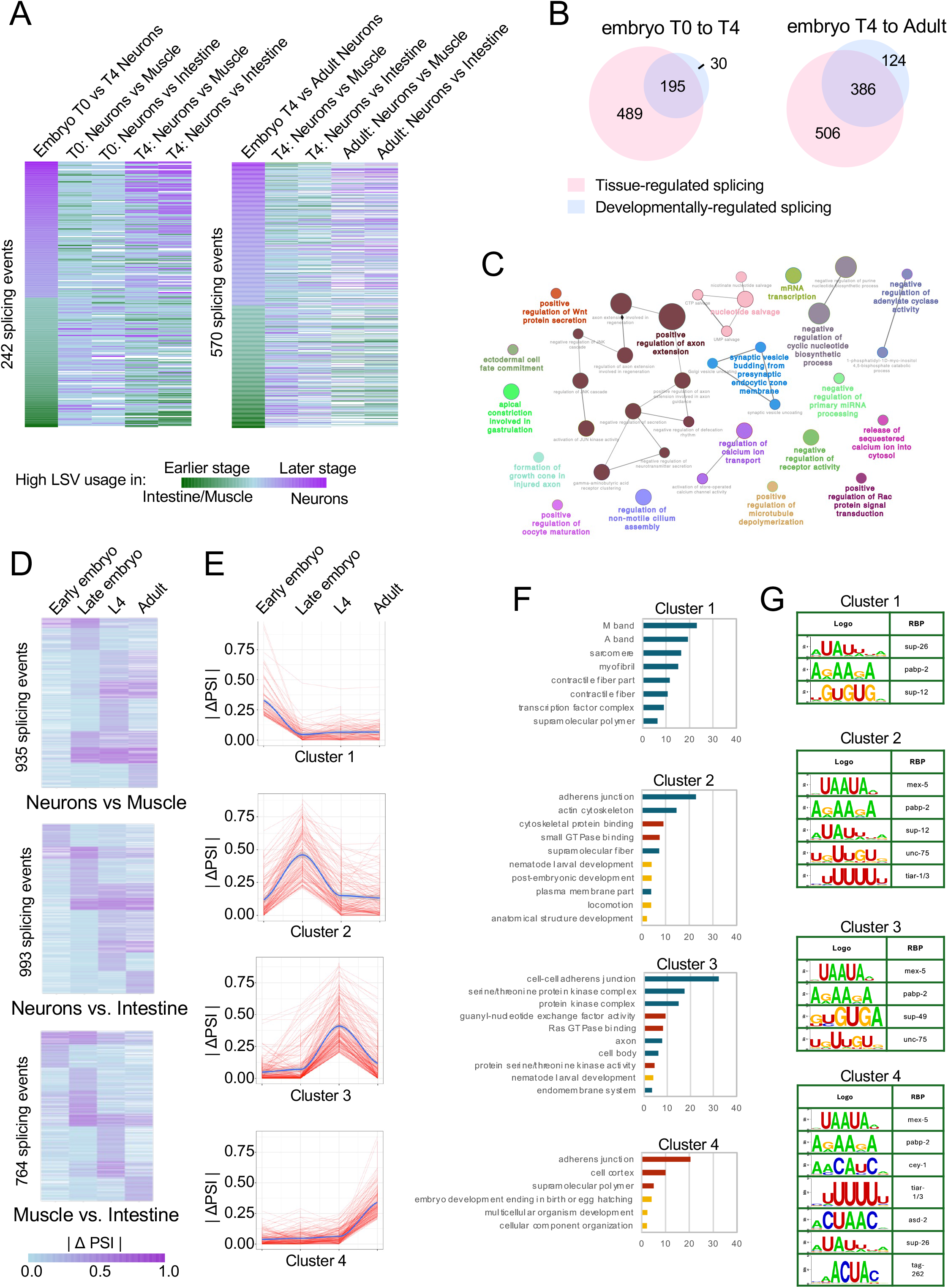
Interplay between developmental- and tissue-specific regulation of splicing programs. A) Left heatmap: 242 alternative splicing events (LSVs) with significant developmental regulation between early and late stages of embryonic neuronal development (Embryo T0 vs. T4 neurons, left column), compared with LSV usage in pairwise tissue comparisons (neurons vs. muscle and neurons vs. intestine at the corresponding developmental stages, T0 and T4, remaining columns). For developmental regulation, green indicates higher LSV usage at T0 and purple at T4; for tissue comparisons, green indicates higher LSV usage in intestine/muscle and purple in neurons. Right heatmap: Comparing LSV usage in 570 developmentally-regulated splicing events between T4 and adult stage neurons with LSV usage values between tissues at the same stages. B) Venn diagrams showing the overlap between genes containing developmentally-regulated splicing events (blue; differences between two time points in neurons) and genes containing tissue-regulated splicing events (pink; differences between neurons and intestine/muscle at the same time points). Left diagram: embryo T0 and T4 comparison. Right diagram: embryo T4 and adult comparison. C) GO term enrichment analysis using ClueGO of developmentally-regulated genes in neurons (all developmental stages combined). D) Heatmaps of tissue-regulated splicing events across development, showing | Δ PSI | for splicing events with significant differences in splicing between at least two stages. Events are grouped by k-means clustering to highlight distinct developmental splicing patterns. E) A set of four splicing trajectories that group tissue-regulated splicing events from all three heatmaps seen in D) that follow a set pattern of | Δ PSI | across development. Blue line is the regression line and the shaded grey area is the 95% confidence interval. Cluster 1 comprises all tissue-regulated splicing events that showed a peak in | Δ PSI | at the early embryonic time point and a subsequent plateau for all subsequent time points. Similarly, Cluster 2 events show a | Δ PSI | peak at the late embryo time point; Cluster 3 events show a | Δ PSI | peak at the L4 stage; and Cluster 4 events show a | Δ PSI | peak at the adult stage. F) GO term enrichment analysis on the events categorized in each cluster from E). Enrichment score bars are colored according to the three GO aspects: blue = Cellular Component, red = Molecular function, yellow = Biological Process. G) MEME-SEA motif enrichment analysis of the tissue-regulated events searched against the CISBP-RNA database gives a list of high-confidence enriched motifs and associated RBPs (p-value <0.01).

We performed GO (Gene Ontology) term enrichment analyses of the functional networks of genes that are D-R in each tissue (all stages combined; **Fig. 2C** and **Supplemental Fig. 2**). Each enrichment analysis reveals biological processes that are relevant to each tissue. For example, GO terms that are directly related to neuronal development, such as *ectodermal cell fate commitment*, and *positive regulation of axon extension*, are present in neuron D-R splicing events (**Fig. 2C**). Similarly, GO terms such as *myoblast fusion*, *actin filament bundle assembly* and *positive regulation of axon extension* are represented in muscle D-R events (**Supplemental Fig. 2C**). Prominent GO terms found in intestinal D-R events were *positive regulation of clathrin-dependent endocytosis/negative regulation of endocytosis*, *regulation of gastrulation* and *embryonic endodermal digestive tract morphogenesis* (**Supplemental Fig. 2F**).

To further dissect the T-R LSVs we identified across development, we next grouped AS events based on any shared splicing trajectories. The absolute difference in isoform usage (|dPSI| of LSV) for every pairwise comparison was plotted as a heatmap clustered by k-means clustering (**Fig. 2D**). In each pairwise comparison, several clusters emerged, grouped based on the developmental stages in which they exhibit the highest or lowest absolute differences in LSV usage. Amalgamating all three pairwise tissue comparisons, we also generated graphs that depict the LSV usage trajectories of these events (**Fig. 2E**). This resulted in ten different splicing trajectories, the first four having peaks of absolute differences at either early embryo, late embryo, L4 or adult (**Fig. 2E**, Clusters 1-4), the next two having a gradual increase or decrease in usage over time (**Supplemental Fig. 3B**, Clusters 5-6), and the remaining four showing peaks and plateaus at various stages (**Supplemental Fig. 3B**, Clusters 7-10).

Given that biologically related networks of splicing events can be instructed by shared splicing mechanisms (Ule & Blencowe, 2019; Van Nostrand et al., 2020), we analyzed each of these ten trajectories for GO term enrichment. GO terms that were frequently enriched in the trajectories were related to cell adhesion and cytoskeletal processes/actin dynamics (**Fig. 2F**, **Supplemental Fig. 3C**), suggesting that these pathways may be significantly impacted by splicing regulation at various stages of development. However, there are instances of trajectory-specific GO terms such as *post-embryonic development* in Cluster 2 with a high differential usage peak at late embryo, and *nematode larval development* in Cluster 3 with a peak at the L4 stage. As the worm progresses to later stages of development, GO terms regarding neuronal development also appear, such as *neurogenesis*, *neuronal cell body*, and *somatodendritic compartment* in Clusters 8 and 10 (**Supplemental Fig. 3C**). These clusters correspond to high differential usage at the late embryo and L4 stages. These splicing trajectories coincide with phases of nervous system development, including the birth of 80 post-embryonic neurons, ventral nerve cord expansion, continuous synaptogenesis, and the synaptic rewiring required to form a mature adult nervous system (Mulcahy et al., 2022; Pathak et al., 2020).

To investigate whether different splicing trajectories were regulated in a coordinated manner by RNA-binding proteins (RBPs), we performed enrichment analysis using MEME-SEA (Bailey & Grant, 2021) and mapped results to known RBP motifs from the CISBP-RNA database (Ray et al., 2013) on clusters 1-4 (**Fig. 2G**). We identified motifs that were significantly enriched in the vicinity of these AS events, recognized by RBPs with known roles in tissue- and/or developmental-specific alternative splicing such as UNC-75 (Kuroyanagi, Watanabe, & Hagiwara, 2013; Norris et al., 2014; Pilaka-Akella et al., 2025), SUP-12 (Anyanful et al., 2004; Ohno et al., 2012), and ASD-2 (Ohno et al., 2008). We also identified over-represented motifs for SUP-26 (Mapes et al., 2010), PABP-2 (Hurschler et al., 2011), MEX-5 (Pagano et al., 2007), and CEY-1 (Arnold et al., 2014), which have known roles in post-transcriptional regulation, RNA localization, stability or translation, but no characterized role in splicing regulation (Peng & Murray, 2025). Some of these RBP-motifs are enriched in multiple clusters, suggesting broad roles for those RBPs in splicing regulation during development. Other enriched motifs are more specific and possibly indicate that a subset of splicing events fall within more specialized networks of co-regulated splicing events.

Finally, we also explored the possibility that groups of splicing events with similar trajectories may also be interconnected within the same protein-protein interaction networks. We conducted a protein-protein interaction analysis using STRING (Szklarczyk et al., 2023) and our observations revealed the presence of interactions among subsets of genes that undergo differential splicing within each cluster. Notably, the most substantial networks were identified within clusters 2-4 that have peaks of differential splicing at the late embryo, L4 and Adult stages (**Supplemental Fig. 3A** and **3C**).

**Figure 3:**
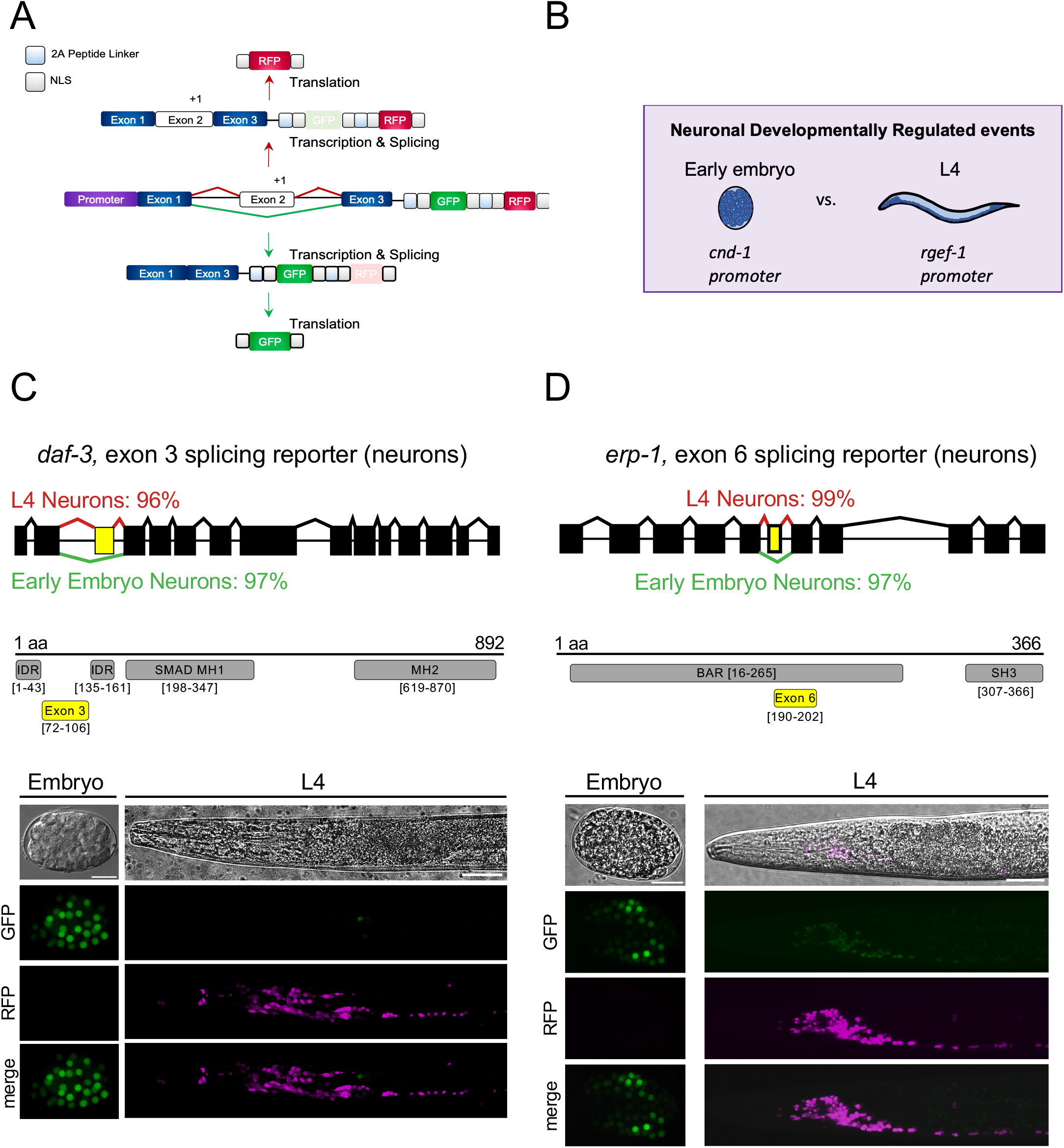
Two-colour reporters of neuron-specific splicing events predicted to be developmentally regulated. A) Two-colour splicing reporter construct consists of a minigene containing an alternatively spliced cassette exon (yellow) with its flanking introns and constitutive exons (blue), cloned upstream of GFP and mCherry (RFP) ORFs. The alternative exon is engineered to include an additional nucleotide (+1), introducing a reading frame shift. Consequently, inclusion of the exon places the RFP in-frame, whereas exon skipping restores the GFP reading frame. Cleavage at 2A peptide sequences and the presence of nuclear localization signals (NLSs) enable the release of fluorescent proteins, which subsequently accumulate within the nucleus. B) Neuron-specific splicing events were examined at two developmental stages. Two-colour splicing reporters driven by the *cnd-1* promoter were used to monitor splicing activity in neuroblasts during early embryogenesis (mid-gastrulation). At the L4 larval stage, splicing was monitored using reporters expressed under the control of the pan-neuronal *rgef-1* promoter, which is expressed in postmitotic neurons. C) Map of the *daf-3*, exon 3 cassette splicing event predicted from RNA-seq data to be predominantly skipped (97%) in early embryonic neuroblasts and then undergo a switch to become highly included (96%) in L4-stage neurons. Diagram below indicates the position of conserved protein domains (grey) and the alternative exon (yellow) with bracketed ranges denoting the corresponding amino acid positions. Fluorescence microscopy of neuronally-expressed splicing reporter for *daf-3*, exon 3 at the embryo (scale bar = 15µm) and L4 stages (scale bar = 30µm). The daf-3 two-colour splicing reporter is expressed under the control of the *cnd-1* promoter in the embryo stage and *rgef-1* promoter for the L4 stage worms. RFP or GFP signals indicate the inclusion or skipping, respectively, of the alternative exon 3. D) Map of the *erp-1*, exon 6 cassette splicing event predicted from RNA-seq data to be predominantly skipped (97%) in early embryonic neuroblasts and then undergo a switch to become highly included (99%) in L4-stage neurons. Diagram below indicates the position of conserved protein domains (grey) and the alternative exon (yellow) with bracketed ranges denoting the corresponding amino acid positions. Fluorescence microscopy of neuronally-expressed splicing reporter for *erp-1*, exon 6 at the embryo (scale bar = 15µm) and L4 stages (scale bar = 30µm). The *erp-1* two-colour splicing reporter is expressed under the control of the *cnd-1* promoter in the embryo stage and *rgef-1* promoter for the L4 stage worms. RFP or GFP signals indicate the inclusion or skipping, respectively, of the alternative exon 6.

Taken together, our results indicate that as tissues differentiate and mature, developmental splicing programs coordinated by upstream regulators help establish tissue identity and function. Our analysis further highlights key biological modules and pathways which are co-regulated via AS.

### Confirmation of tissue- and developmentally-regulated splicing events in vivo via two-colour reporters

To validate our analysis of D-R and T-R splicing events, we selected several alternatively spliced exons that exhibit switch-like behaviour with high levels of inclusion in one condition and low levels of inclusion in another condition (|dPSI|>=0.5). Generally, these exons also exhibit a high degree of conservation and play a pivotal role in influencing the conditions in which they are regulated (Barbosa-Morais et al., 2012; Bush et al., 2017; Koterniak et al., 2020; Mazin et al., 2021; Xing & Lee, 2005). We selected two D-R regulated events (*daf-3*, exon 3 and *erp-1*, exon 6) and one T-R regulated event (*kvs-5*, exon 6) to validate using a two-colour splicing reporter system that enables the *in vivo* visualization of alternative splicing patterns at single-cell resolution using fluorescence microscopy (Calarco & Pilaka-Akella, 2022; Norris et al., 2014). Each reporter contains a minigene comprising the alternative exon of interest along with its flanking introns and exons (**Fig. 3A**). The minigene is placed upstream of tandem GFP and RFP ORFs that are encoded in different reading frames. The sequence of the alternative exon was modified to shift the reading frame such that exon skipping yields GFP expression, whereas exon inclusion (+1 frameshift) results in RFP expression. A tissue-/developmental-stage specific promoter positioned upstream of the minigene drives the expression of the reporter. 2A peptide sequences and nuclear localization signals were placed upstream and downstream of each fluorescent protein, respectively (Ahier & Jarriault, 2014; Lyssenko et al., 2007). This results in the cleavage and concentration of fluorescent proteins to the nucleus, enhancing resolution.

The switch-like exons from the *daf-3* (exon 3) and *erp-1* (exon 6) genes were identified as D-R events from the RNA-seq analysis to undergo differential inclusion between early embryonic and L4 neuronal stages. Two-colour reporters were made with *cnd-1* and *rgef-1* promoters, driving expression in embryonic neuroblasts and mature neurons, respectively (**Fig. 3B**; (Hallam et al. 2000; Altun-Gultekin et al. 2001; Chen et al. 2011). The *daf-3* gene encodes a Smad protein that is expressed in neurons as well as many other tissues and positively regulates larval dauer formation as part of a TGF-β-related signaling pathway (Patterson et al., 1997). Exon 3 of *daf-3* falls squarely between two intrinsically disordered regions (IDRs) and was measured from our RNA-seq analysis to be predominantly skipped in early embryos, with a developmental shift to being mostly included at the L4 stage (**Fig. 3C**, top). Confocal imaging of the two-color reporters confirmed this pattern, with the almost exclusive expression of the GFP (skipped) isoform in embryos and the RFP (included) isoform in L4 animals (**Fig. 3C**, bottom). The gene *erp-1* encodes an endophilin related protein involved in synaptic vesicle biology (S. C. Yu et al., 2018) and the alternatively spliced exon 6 overlaps with an N-terminal BAR-domain (**Fig. 3D**). Exon 6 of *erp-1* was confirmed to be more skipped in the early embryo and more included at the L4 stage. Lastly, the *kvs-5* gene encodes a potassium voltage-gated channel subunit and exon 6 was found in our dataset to demonstrate switch-like behaviour between L4 stage neurons and body-wall muscle (**Supplemental Fig. 4**). This exon is situated between sequences encoding potassium channel domains and demonstrates switch-like behaviour between L4 stage neuron and muscle tissues. To observe this event, we generated two reporters using the *rgef-1* and *myo-3* promoters to drive expression in neurons and body-wall muscle (Okkema et al., 1993), respectively (**Supplemental Fig. 4**). Both reporters were co-injected into the same animal, enabling simultaneous visualization of splicing patterns in the two tissues. Confocal imaging shows almost exclusively RFP (exon included) signal in muscle nuclei and predominantly GFP (exon skipped) signal in neurons (**Supplemental Fig. 4C**). Taken together, our two-colour reporters independently validate and support our differential splicing transcriptome analysis.

### Identification of a more expansive catalog of regulated microexons in the C. elegans genome

Microexons are a class of exons spanning 3 to 27 (or in some studies, up to 51) nucleotides and require specialized mechanisms for inclusion due to their small size. In mammals, microexon inclusion is critical for the proper development and function of the nervous system (Irimia et al., 2014b). Previous work identified a set of 75 microexons in the *C. elegans* genome (Weinreb et al., 2025). However, curation of these exons has relied on existing EST/cDNA coverage and whole animal transcriptome data. Leveraging our tissue- and stage-specific transcriptome and translatome datasets (see Methods), we sought to more comprehensively identify microexons within the *C. elegans* genome.

Given their small size, microexons are frequently missed by conventional RNA-seq mapping tools. We adapted a pipeline from (H. Yu et al., 2022) to improve microexon discovery (**Fig. 4A**). The pipeline involves the joint usage of STAR (Dobin et al., 2013), a standard mapping tool, and OLego (Wu et al., 2013), which uses small alignment seeds for *de novo* mapping of spliced RNA-seq reads. Junctions detected by both were combined for downstream STAR mapping. Output files were used to generate a new GTF file containing transcript information and fed into MAJIQ (Vaquero-Garcia et al., 2016) for alternative splicing analysis. A list of putative microexons was compiled and filtered to remove certain classes of exons including: 1) first exons, 2) exons having coordinates coinciding with existing longer exons and 3) exons with insufficient read support. This first, lower stringency list contained 176 microexons (**Fig. 4B; Supplementary Table S3**), substantially exceeding the 24 microexons detected previously using standard mapping approaches in our previous analysis of L4 tissue transcriptomes (Koterniak et al., 2020). We examined this list for regulatory features such as dinucleotide splice site sequences, frame preservation and conservation. We determined that 143 microexons were flanked by canonical donor and acceptor splice sites (5’ GT and 3’ AG), and the remaining 33 were flanked by other splice site dinucleotide sequences. Notably, within the former group of microexons with canonical splice sites, a substantial majority (85%) are a multiple of three and maintain the reading frame when included in transcripts. Conversely, in the latter group of microexons with non-canonical splice sites, only 55% of these microexons preserve the reading frame. Lastly, we determined a stringent list of microexons with both conserved splice sites and conserved microexons (n=100) and found that the majority of these microexons are both frame preserving and feature canonical splice sites (**Supplementary Table S4**).

**Figure 4:**
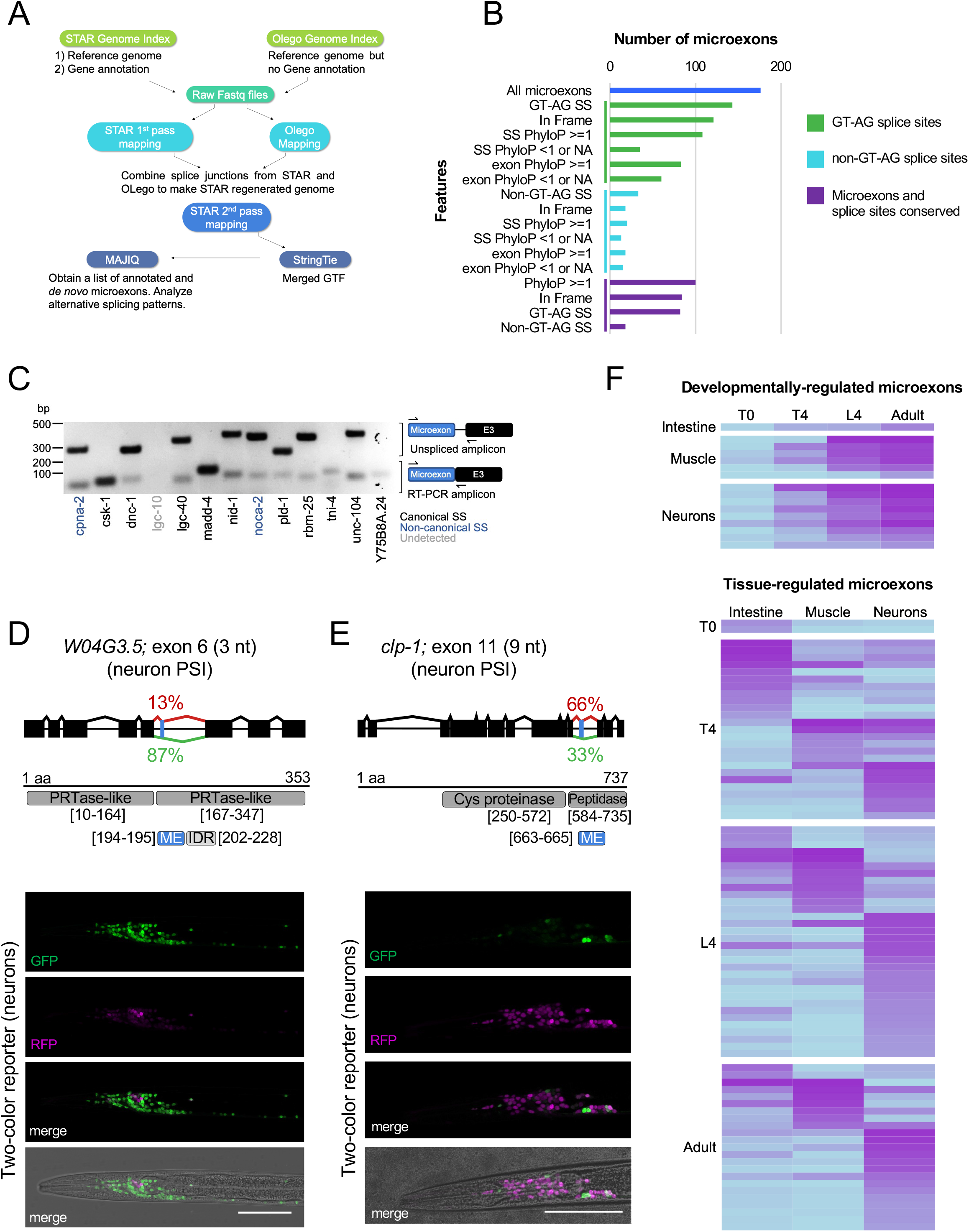
Network and splicing trajectories of tissue-specific splicing events across development. A) Overview of the computational RNA-seq alignment pipeline designed to improve microexon detection. Adapted from Yu et al. (2022). B) Breakdown of detected microexons according to splice site type (canonical GT-AG vs. non-canonical), frame preservation, and evolutionary conservation (PhyloP scores) of splice sites (SS) and microexons. C) RT-PCRs on 13 randomly selected microexons not annotated in current genome annotation, but detected in microexon pipeline described in A). RT-PCR primers were designed to amplify a sequence between ∼70-120bp in length spanning the microexon and downstream exon (E3) and correspond to the lower bands. The genes containing the microexons are coloured according to the microexon having canonical 5ʹ and 3ʹ splice sites (black, n=10), non-canonical (blue, n=2) and putative *de novo* microexons that could not be detected by RT-PCR (grey, n=1). Upper bands were Sanger sequenced and found to be unspliced amplicons with retained introns (data not shown). D) Top: Map of the *W04G3.5*, exon 6 microexon (3nt) splicing event predicted from RNA-seq data to be 87% skipped and 13% included in neurons. Diagram below indicates the position of conserved protein domains (grey) and the microexon (ME; blue) with bracketed ranges denoting the corresponding amino acid positions. Bottom: Fluorescence microscopy of a worm expressing the *W04G3.5*, exon 6 two-colour splicing reporter under the control of the *rgef-1* pan-neuronal promoter. RFP or GFP signals indicate the inclusion or skipping, respectively, of the alternative microexon. Scale bar = 50µm. E) Top: Map of the *clp-1*, exon 11 microexon (9nt) splicing event predicted from RNA-seq data to be 33% skipped and 66% included in neurons. Diagram below indicates the position of conserved protein domains and the microexon as described in E). Bottom: *clp-1*, exon 11 two-colour splicing reporter under the control of the *rgef-1* pan-neuronal promoter. RFP or GFP signals indicate the inclusion or skipping, respectively, of the alternative microexon. Scale bar = 50µm. F) Top: Heatmaps of developmentally-regulated microexons in intestine, muscle, and neuronal tissues, showing progressive inclusion from T0 (early embryo) through T4 (late embryo), L4, and adult stages (total # microexons = 16). Microexons were required to demonstrate a ≥30% increase in inclusion between T0 and adult stages and stepwise increases across each successive stage. Bottom: Heatmaps of tissue-regulated microexons that demonstrate a ≥30% difference in PSI value between at least two tissues at each developmental stage (total # microexons = 82).

In both the broad and stringent microexon lists, we noted that a substantial portion of microexons were previously non-annotated (40% and 42%, respectively). We tested the expression of a set of 13 high-confidence unannotated microexons through RT-PCRs using total RNA from N2 animals. All of these microexons ranged from 6-27nt in length and 2/13 had non-canonical splice sites. Primers were designed to amplify a fragment spanning the microexon and the 5’ end of the downstream exon. Out of the 13 microexons, the expression of 12 was validated using our RT-PCR approach (**Fig. 4C**), underscoring the effectiveness of the computational strategies employed in our unannotated microexon discovery.

GO analysis suggests that this set of 100 high confidence microexons primarily regulates neuronal activity, specifically synaptic biology, axon guidance and potassium channel activity (**Supplemental Fig. 5A**). This observation aligns with studies across human, mouse, *D. melanogaster*, and *C. elegans*, which also show that genes containing microexons are enriched for neuron-related biological processes (Choudhary et al., 2021; Irimia et al., 2014b; Pang et al., 2021; Parada et al., 2020). Additionally, genes with microexons are enriched for roles in muscle development, mating behaviour, and in splicing and AS. Given the observation that microexon-containing genes are frequently linked to neuronal processes, we focused on alternatively spliced microexons (PSI <=0.95) identified in our dataset that were predicted to generate both included and skipped isoforms in neurons. Using the two-color splicing reporter system described above, we visualized the splicing patterns of *W04G3.5* exon 6, *clp-1* exon 11, *kvs-5* exon 6, and *madd-4* exon 6 (**Figs. 4D** and **E**, **Supplemental Fig. 4**, **Fig. 5**). We validated that all these microexons were included in the worm nervous system, generally recapitulate the expected splicing patterns measured from our transcriptome analysis, and in several cases, displayed neuron subtype-specific splicing patterns.

**Figure 5:**
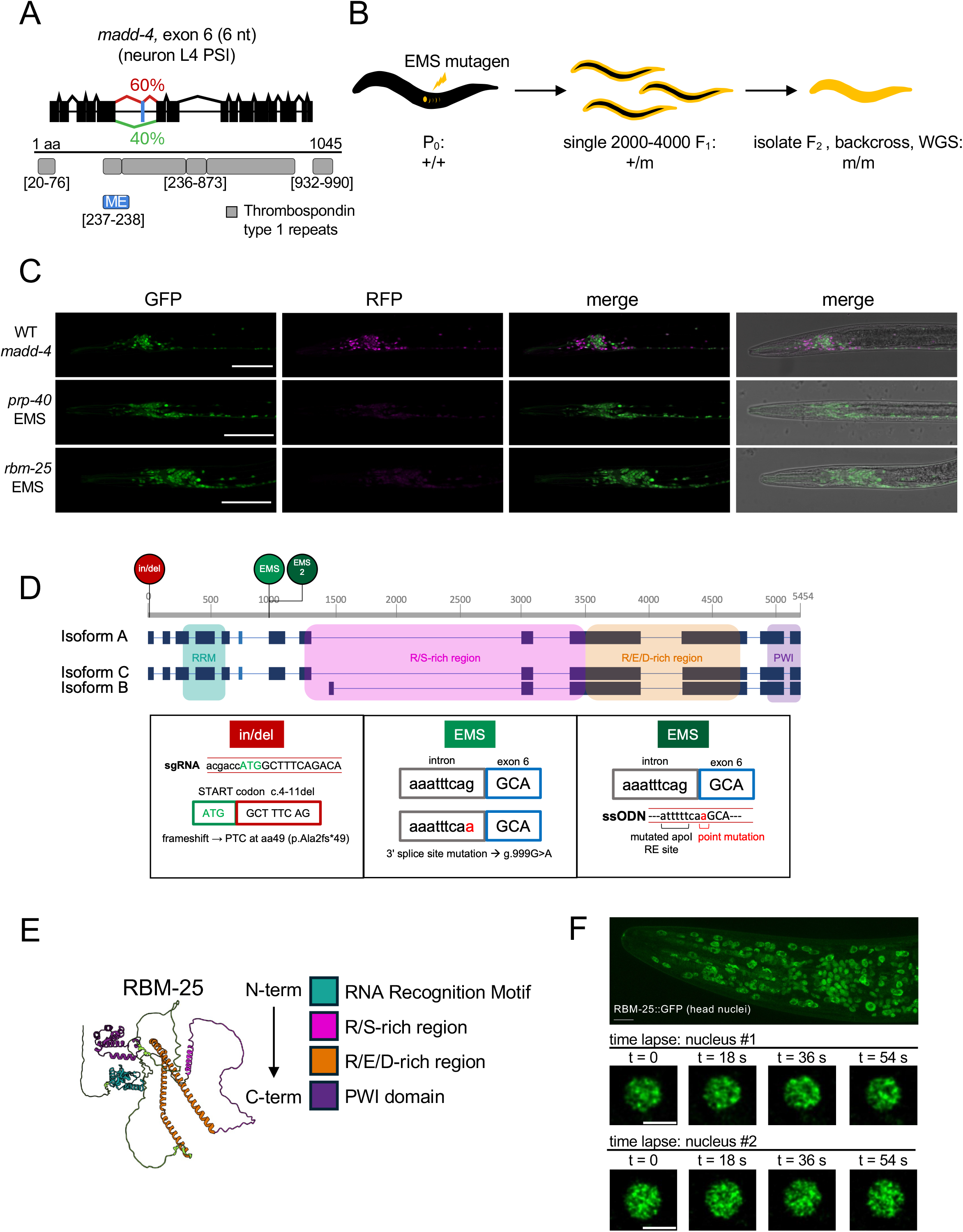
Network and splicing trajectories of tissue-specific splicing events across development. A) Map of the *madd-4*, exon 6 microexon (6nt) splicing event predicted from RNA-seq data to be 40% skipped and 60% included in neurons. Diagram below indicates the position of conserved protein domains (grey) and the microexon (ME; blue) with bracketed ranges denoting the corresponding amino acid positions. B) The *madd-4*, exon 6 two-colour reporter was stably integrated into the genome. A forward genetics screen was conducted using ethyl methanesulfonate (EMS) and approximately 2000 F1 worms (4000 haploid genomes) were singled and cultured for mutation screening in the F2 generation. F2 worms that demonstrated a perturbation splicing reporter fluorescence pattern were backcrossed and selected for whole-genome sequencing to determine causal mutations. C) Fluoresence microscopy of the *madd-4*, exon 6 microexon two-colour reporter under wild-type and mutant conditions. Shown from left to right: the wild-type madd-4 exon 6 microexon reporter, which was utilized in an EMS forward genetic screen to identify regulators of microexon inclusion. Among the mutants recovered from this screen were alleles of *prp-40* and *rbm-25*. D) Schematic representation of *rbm-25* isoforms and mutations analyzed in this study. Diagram showing the *rbm-25* gene structure, three annotated transcript isoforms (A–C) and conserved proteins domains, RRM (RNA recognition motif) and PWI domains are highlighted in purple and green, respectively. A putative *de novo* microexon is coloured in blue. The CRISPR-Cas9 generated in/del mutation (red) targets the start codon, resulting in a c.4–11 deletion that causes a frameshift and PTC at amino acid 49 (p.Ala2fs*49). The EMS mutant (green) carries a single nucleotide substitution (g.999G>A) that disrupts the 3ʹ splice site of exon 6. A second EMS allele was generated using CRISPR–Cas9 and a single-stranded oligonucleotide donor (ssODN) to recreate the same point mutation and assess its impact on splicing. E) AlphaFold prediction of the RBM-25 protein in *C. elegans*. The two conserved domains and other regions are listed in order of N-term to C-term. There are also three additional unstructured regions, two flanking the RRM and another between the R/E/D-rich region and the PWI domain. The N-term RRM is expected to form a four-stranded antiparallel β-sheet and two α-helices, while the C-term PWI domain forms a four-helix bundle structure. F) Top: Fluorescence microscopy of a worm expressing GFP-tagged RBM-25 shows subcellular localization in the head nuclei. Scale bar: 10µm? The lower panels show two individual nuclei imaged over 54 seconds. Scale bars: 5 µm.

We sought to determine what fraction of microexons are differentially alternatively spliced and to characterize the range of their inclusion levels. We plotted the maximum difference in PSI values between any two samples for each microexon and observed a dynamic spectrum of inclusion levels (**Supplemental Fig. 5B**). We found that 35% of microexons have a switch in PSI value of >=0.5 and 9% have an extreme switch in PSI value of >=0.9 between any two conditions. To visualize microexon splicing dynamics in more detail across the intestine, body-wall muscle, and neurons throughout development, we identified developmental-or tissue-regulated microexons that had >=0.3 difference in PSI and generated PSI value heatmaps (**Fig. 4F**). We found 16 microexons with differing inclusion levels during development, with 9 of these splicing events occurring in neurons. We also found 82 tissue-biased microexons (**Fig. 4F**). Notably, during embryonic stages, the intestine shows the highest number of differentially included microexons. However, beginning at the L4 stage, this pattern shifts, with neurons exhibiting the greatest number of included microexons.

Finally, we also performed a motif enrichment analysis to test whether any RNA-binding protein motifs were over-represented in the introns flanking microexons (**Supplemental Fig. 5C**). This analysis revealed an enrichment in EXC-7, ASD-1/FOX-1, and SAP-49 motifs in the introns flanking microexons. Positional distribution of motifs along the length of sequences centered at the microexon show that all four RBPs had the most significant peaks of motif enrichment downstream of the 5’ splice site. Taken together, our results have expanded the number of microexons in the *C. elegans* genome and reveal conservation and regulatory patterns suggesting their functional importance in shaping tissue identity and function.

### A mutagenesis screen identifies rbm-25 as a regulator of microexon inclusion

To discover regulators of microexon inclusion in *C. elegans*, we undertook an unbiased, forward mutagenesis screen. We first selected a two-colour microexon splicing reporter for the gene, *madd-4*, an ortholog of the mammalian *punctin-1* and *punctin-2* involved in axon guidance and synapse organization (Kratsios et al., 2015; Seetharaman et al., 2011)(**Fig. 5A**). The reporter minigene contained exon 6, a six nucleotide microexon, within a Thrombospondin type-1 (TSP1) repeat that showed robust splicing in neurons, consistent with our RNA-seq data (60% exon inclusion, **Fig. 5A** and **5C**, left). Animals expressing these reporters were mutagenized with ethyl methanesulfonate (EMS) and approximately 2,000 F1 progeny were clonally isolated (∼4,000 haploid genomes) and cultured for screening in the F2 generation (**Fig. 5B**). Our screen recovered 5 mutant alleles in total. After performing whole genome sequencing and comparing mutant with non-mutagenized reporter animal genomes, we identified likely causal mutations for three of these recovered mutants. Importantly, we isolated two alleles of *prp-40*, which was also recently demonstrated to regulate microexon inclusion (Choudhary et al., 2021), thereby validating the ability of our screen to identify bona fide regulators of microexon splicing. Intriguingly, we also isolated an allele of *rbm-25*, a gene linked to alternative splicing but not yet investigated in the context of microexon splicing in any organism.

RBM-25 is an RNA-binding protein that is conserved across eukaryotes. In many species, including humans and worms, RBM-25 (also referred to as RED120 in mammals, and Snu71 in yeast), contains an N-term RRM (RNA Recognition Motif) domain, a central R/E/D-rich region and a C-term PWI (Proline-Tryptophan-Isoleucine) domain (**Fig. 5D and 5E**; Fortes 2007; Carlson 2017). Orthologous RBM-25 protein sequences were retrieved from OrthoDB (Zdobnov et al., 2021) and aligned across 14 eukaryotic model organisms (**Supplemental Fig. 6)**. Phylogenetic analysis revealed that *Saccharomyces cerevisiae*, *Schizosaccharomyces pombe*, and *Arabidopsis thaliana* form independent branches relative to the metazoan RBM-25 proteins. Sequence alignments show a proline-rich region at the N-term that is well-defined in *X. laevis*, *G. gallus*, and mammals, but is somewhat less noticeable in *C. elegans*, consistent with previous analysis (Fortes et al., 2007). In worms, current gene models suggest *rbm-25* encodes three annotated isoforms (A-C). Isoforms A and C arise from the use of an alternative 3’ splice site in exon 10 and differ by only two amino acids, with isoform C containing the additional residues. Isoform B is a truncated isoform lacking the first seven exons of the longer isoforms and the encoded RRM domain entirely (**Fig. 5D**). However, our RNA-seq analysis identifies junction-spanning reads connecting exon 7 of isoforms A/C to the first exon of isoform B, suggesting that the current isoform annotations may be incorrect (data not shown). We also discovered an unannotated microexon in *rbm-25* downstream of the exons encoding the RRM and validated its inclusion among our aforementioned RT-PCR validations (**Fig. 4C**). The recovered *rbm-25* EMS mutant harbors a single nucleotide substitution (g.999G>A) that disrupts the 3ʹ splice site upstream of exon 6 (**Fig. 5D**). To validate the sufficiency of this point mutation in causing defects in microexon splicing, we used CRISPR/Cas9 genome editing to recreate this single-base edit (Eroglu et al., 2023; Paix et al., 2016). Confocal microscopy of this new strain recapitulated the splicing pattern observed in the EMS mutant (**Supplemental Fig. 7**).

In humans, the RRM domain and R/E/D-rich regions mediate many of the interactions between RBM25 and other splicing factors or binding targets (Carlson et al., 2017; J. D. Li et al., 2024; Sénéchal et al., 2023; Zhou et al., 2008). Since the EMS point mutation lies in exon 6 of *rbm-25*, which is located downstream of the exons encoding the N-terminal RRM (RNA-recognition motif) domain (**Fig. 5D**), we also sought to generate an independent mutant allele of *rbm-25*. Therefore, we targeted the start codon with CRISPR/Cas9 and a non-homologous end joining (NHEJ) repair event produced an 8bp deletion immediately downstream of the ATG, causing a frameshift and introducing a premature termination codon (PTC) at amino acid position 49. Confocal microscopy of the two-colour reporter spicing pattern confirmed this mutant strain also recapitulates the EMS mutant phenotype of decreased *madd-4* microexon inclusion (**Supplemental Fig. 7)**.

Consistent with a role in alternative splicing, expression of endogenously tagged RBM-25::GFP in the worm showed largely nuclear localization (**Fig. 5F**). GFP signal appeared ubiquitous across most cell types, identified based on position and morpohology, suggesting that RBM-25 is expressed broadly. Live imaging of individual nuclei over an ∼1 minute time frame showed that GFP signal was enriched in discrete, dynamic nuclear puncta rather than diffusely throughout the nucleus. Collectively, our analysis of several mutant alleles, as well as the subcellular localization pattern of RBM-25::GFP, strongly suggests that *rbm-25* is an activator of microexon splicing.

### rbm-25 mutants exhibit multiple physiological and behavioral phenotypes

We first assessed broad phenotypes of *rbm-25* mutants with assays measuring reproductive fitness and lifespan. Measurements of brood size between N2 wild-type and *rbm-25* showed that mutants have 26% less progeny (mean brood size: 324 +/-59.8 for N2 wild type and 241 +/-43.1 for *rbm-25*; Wilcoxon rank-sum test, p = 1.3e-08; **Fig. 6A**). This is especially noticeable during the first three days of egg-laying and hatching, when the rate of progeny being laid per worm is reduced in the mutants (**Supplemental Fig. 8A**). We also found that while N2 wild-type animals have a median lifespan of 12 days post-adulthood, *rbm-25* mutant animals have median lifespan of 4 days (mean survival: 11.7 days (SE = 0.453) for N2 and 6.8 days (SE = 0.425) for *rbm-25*; log-rank test, p = 3e-11; **Fig. 6C**). These results suggest that defects in RBM-25 activity can lead to broad phenotypes affecting lifespan and reproductive fitness, which may include germline proliferation, oocyte and/or sperm production, or egg laying.

**Figure 6:**
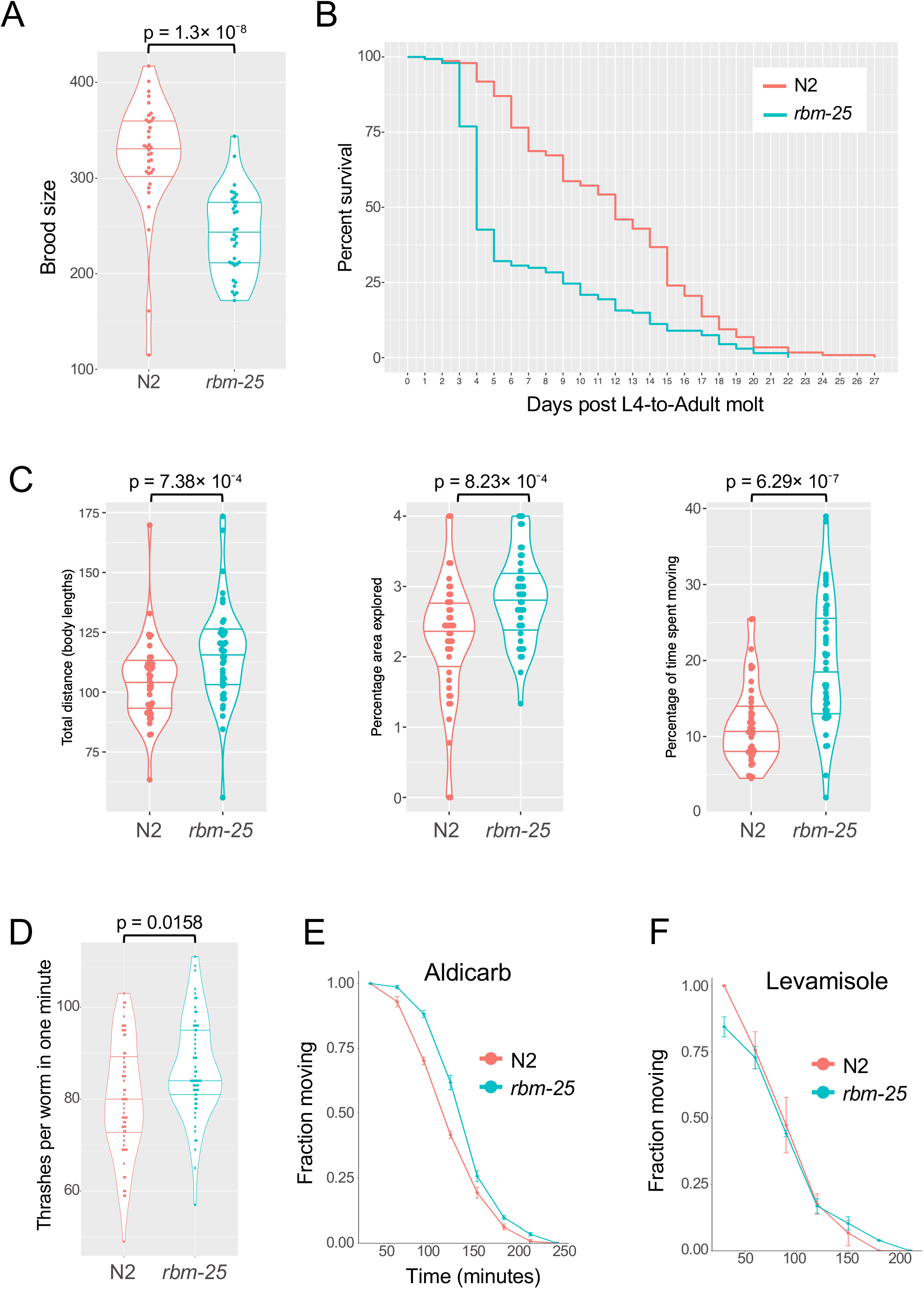
Fitness, lifespan, locomotory and pharmacological responses in r*bm-25* mutants. A) Brood size assay measuring the total number of viable progeny and unhatched eggs laid by N2 wild-type and *rbm-25* mutant worms. Mean brood size was 324 (SD = 59.8) for N2 wild type and 241 (SD = 43.1) for *rbm-25*. Differences between genotypes assessed using Wilcoxon rank-sum test. p-value=1.3e-08. Assays were performed in duplicates with 16-20 animals per strain, for a total of n = 35 per strain after outlier removal. B) Lifespan assay of N2 and *rbm-25* worms using Kaplan–Meier survival curves. Mean survival was 11.7 days (SE = 0.453) for N2 and 6.8 days (SE = 0.425) for *rbm-25*. Survival differences between genotypes was assessed by a log-rank test. p= 3e-11. Assay started with n = 150 animals per strain. Lost worms were censored from the final count, which was n = 132 for N2 and n = 142 for *rbm-25*. C) Measurement of motor differences between N2 wild-type and *rbm-25* mutant worms using locomotion assays on solid agar. Comparisons between genotypes performed using Wilcoxon rank-sum test. n = 44 for each strain. Left: Total distance travelled in body lengths for a duration of 10 minutes. Mean normalized distance was 104.4 body lengths (SD = 16.7) for N2 and 116.1 body lengths (SD = 20.6) for *rbm-25.* Middle: Mean percentage of grid area explored was 2.27 (SD = 0.84%) for N2 and 2.84% of possible boxes (SD = 0.63%) for *rbm-25*. Right: Mean percentage of time spent moving was 11.46% (SD = 5.18%) for N2 and 19.47% (SD = 8.43%) for *rbm-25*. D) Thrashing assay to measure locomotory differences between N2 wild-type and *rbm-25* mutant worms in liquid medium. Mean number of thrashes in one minute was 80.2 (SD = 12.4) for N2 and 86.8 (SD = 10.8) for *rbm-25.* Pairwise comparison between strains using a Welch’s two sample t-test. p=0.0158. n = 28 for N2 and n = 30 for *rbm-25*. E) Aldicarb response curves measuring paralysis for N2 wild-type and *rbm-25* mutant worms. Time-to-paralysis was analyzed using Kaplan–Meier survival curves, with differences between genotypes assessed by a log-rank test. p-value=0.001. Assay performed in triplicates with 20-30 worms per plate. F) Levamisole response curves for N2 wild-type and *rbm-25* mutant worms. No significant difference in response between genotypes. Assay performed in triplicates with 25-30 worms per plate.

Given the enrichment of microexons within genes associated with neuromuscular development and physiology, we also sought to determine whether *rbm-25* mutants exhibit defects in locomotion and synaptic signalling. Using a live imaging system and automated tracking software (see Methods for details), we quantified a variety of behavioural parameters (**Fig. 6C** and **Supplemental Fig. 8**). *rbm-25* mutants exhibited significantly increased total distance traveled, plate area exploration, and percentage of time spent moving compared with wild-type animals on solid medium (**Fig. 6C**). In liquid medium, *rbm-25* mutants showed a significant increase in thrashing rate compared to wild-type animals (mean # thrashes in one minute: 80.2 +/-12.4 for N2 and 86.8 +/-10.8 for *rbm-25*; Welch’s two-sample t-test, p = 0.0158; **Fig. 6D**).

We also examined the integrity of motor neuron circuits that direct locomotion using pharmacological assays probing motor neuron synaptic signalling to postsynaptic muscle cells. The drug aldicarb inhibits acetylcholinesterase, resulting in a buildup of acetylcholine at the neuromuscular synaptic cleft that causes muscle hypercontraction and subsequent paralysis (Mahoney et al., 2006). Resistance or hypersensitivity to aldicarb can indicate defects in neuromuscular synaptic transmission, as mutants with reduced presynaptic acetylcholine release exhibit delayed paralysis, whereas mutants with increased acetylcholine release paralyze more rapidly. In conjunction with the aldicarb assay, we also performed a levamisole assay. Levamisole acts as an agonist of nicotinic acetylcholine receptors on muscle and can be used to determine whether perturbations in cholinergic neurotransmission arise from presynaptic or postsynaptic defects (Davis & Tanis, 2022). Levamisole sensitivity can indicate altered post-synaptic receptor function, while resistance can indicate defects in the receptor or muscle response. *rbm-25* mutants exhibited significantly greater resistance to aldicarb-induced paralysis, with a mean time to 50% paralysis of 136.7 min (SD = 2.5 min) compared with 111.8 min (SD = 2.2 min) in wild-type worms (Kaplan–Meier analysis; log-rank test, p = 0.001; **Fig. 6E**). However, no significant difference in paralysis kinetics was observed between wild-type and *rbm-25* following levamisole treatment (**Fig. 6F**). Taken together, these results suggest that reduced RBM-25 activity can lead to presynaptic defects in motor neurons, and these defects may contribute to altered animal behaviour observed in mutants. Additionally, our data indicate that RBM-25 plays a key role in the role of animal fertility, longevity, and behaviour.

### RBM-25 preferentially regulates microexons and forms a complex with PRP-40

To further validate our *rbm-25* mutant alleles, we examined the splicing of three microexons in our *rbm-25* CRISPR-generated allele backgrounds, using bulk cDNA sequencing. First, we designed RT-PCR primers that would amplify sequence from within the upstream to downstream exons (similar to the schema presented in **Fig. 4C**). The resulting amplicons were subjected to nanopore sequencing to identify all the splice isoforms present and to quantify their relative abundances. For each microexon, inclusion in mature transcripts was significantly reduced in both mutant alleles compared to the reference control strain (**Fig. 7A; see Methods**).

**Figure 7:**
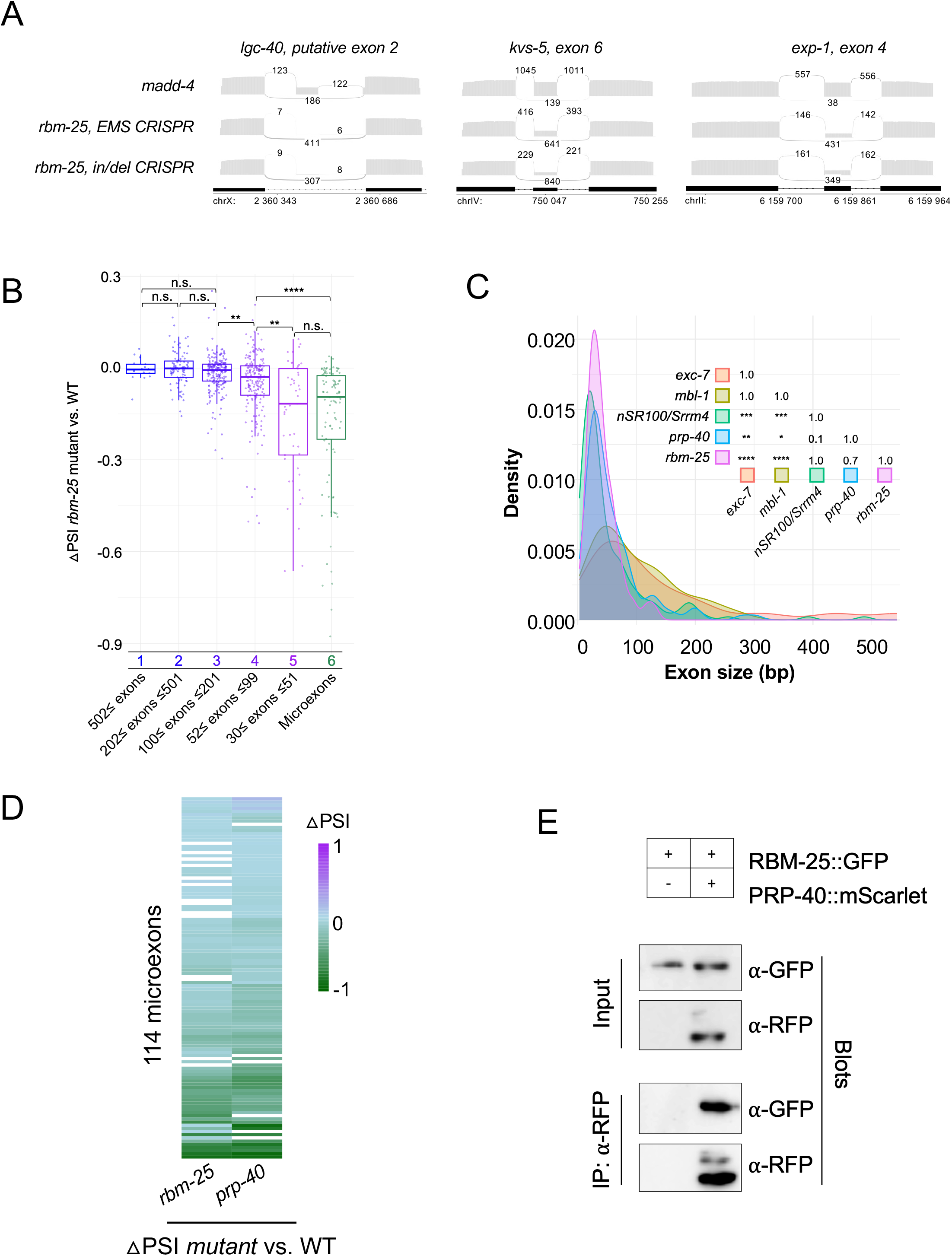
Size-preferential activity of RBM-25 for small exons and co-regulation of microexon inclusion with PRP-40. A) Batch RNA-sequencing of three example microexons of the genes *lgc-40*, *kvs-5* and *exp-1* in the *madd-4* (wild-type), *rbm-25* EMS CRISPR and *rbm-25* in/del CRISPR strains. Sequencing was performed on RT-PCR samples generated using primers complimentary to the upstream and downstream exons flanking the microexons. Sashimi plots show junctions usage and read counts corresponding to microexon-included and skipped reads. Position and coordinates of exons are denoted. B) Distribution of ΔPSI values for all alternatively spliced cassette exons, indicating changes in exon inclusion levels between *madd-4* wild-type and *rbm-25* mutant samples. Each dot represents an individual exon and exons are grouped by size. A ΔPSI value of <0 indicates decreased inclusion of the exon in the *rbm-25* mutant. Pairwise comparisons between conditions were performed using the Wilcoxon rank-sum test and p-values were adjusted for multiple comparisons using the Bonferroni correction. P-value of pairwise comparison between bin 4 (52≤ exons ≤99) and bin 6 (microexons) = 1.7 x 10^-7^. C) Distribution of exon size for cassette exons regulated in five mutants: *rbm-25* (pink), *prp-40* (blue, Choudhary et al. 2021) and mouse *nSR100/Srrm4* (green, Quesnel-Vallières et al. 2015), *mbl-1* (yellow) and *exc-7* (orange; Napier-Jameson et al. 2024). RNA-seq processing for each mutant involved a targeted search for microexons. Regulated exons plotted demonstrated and significant change in inclusion between wild-type and mutant conditions. Pairwise comparisons between conditions were performed using the Wilcoxon rank-sum test and p-values were adjusted for multiple comparisons using the Bonferroni correction. P-values are shown in matrix (* ≤ 0.05, ** ≤ 0.01, *** ≤ 0.001, **** ≤ 0.0001). D) Heatmap of the ΔPSI values of all microexons between and each mutant and WT conditions (*rbm-25* vs. WT and *prp-40* vs. WT). ΔPSI ranges from 1 (purple; more inclusion in WT) to -1 (green; more inclusion in mutant). E) Lysates expressing RBM-25::GFP or co-expressing RBM-25::GFP and PRP-40::mScarlet were extracted. Input blots using anti-GFP antibody (α-GFP) and anti-RFP antibody (α-RFP) confirm the presence of RBM-25::GFP and PRP-40::mScarlet tagged proteins, respectively, consistent with the expression table above for the two lysates. Lysates co-expressing RBM-25::GFP and PRP-40::mScarlet were immunoprecipitated with α-RFP and analyzed by Western blotting with α-GFP and α-RFP antibodies.

**Figure 8:**
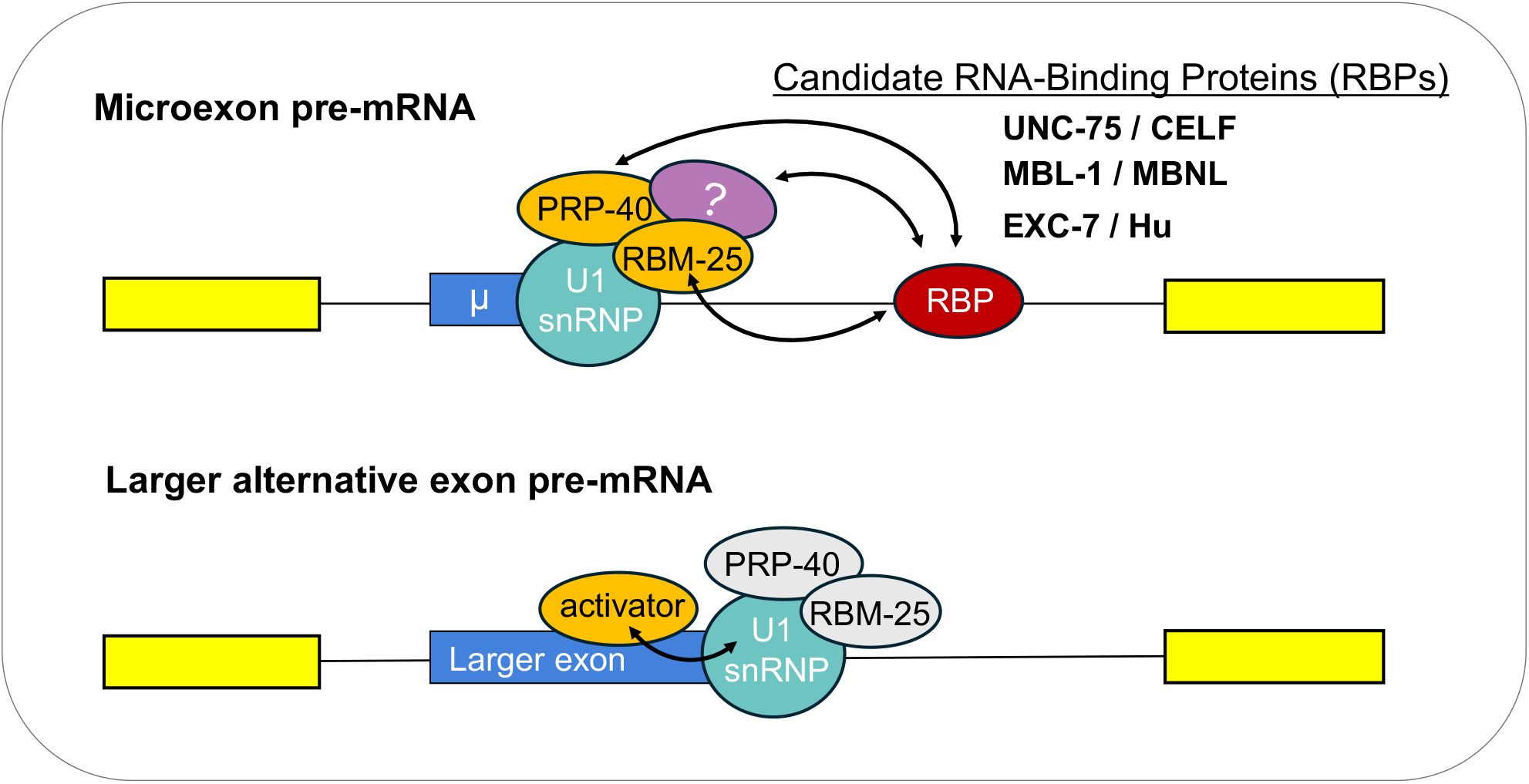
Model for RBM-25, PRP-40, and RBP-mediated microexon inclusion. Top image) To serve as a central regulator of microexon inclusion in multiple tissues, RBM-25 and its interaction partner PRP-40 (orange), and the U1 snRNP (light blue) are likely recruited to microexon (blue) 5′ splice sites via interactions with different tissue-specific RNA-binding proteins (RBPs) such as UNC-75, EXC-7, and MBL-1, and co-regulators (red and purple ovals). Bottom image) For longer alternative exons, PRP-40 and RBM-25 (grey) are more dispensable, likely due to the availability of other activators that bind to exonic splicing enhancer sequences and stimulate U1 snRNP recruitment.

We next asked whether RBM-25 is specifically required for microexon inclusion or instead functions as a more general regulator of splicing. We performed RNA-seq on *rbm-25* in/del mutants and analyzed the data using our microexon discovery pipeline to enhance the detection of microexons while also identifying RBM-25-regulated splicing events. We selected cassette exons that had read support in both a reference strain (expressing the *madd-4* two colour reporter) and *rbm-25* mutants, divided them into bins based on exon size and plotted ΔPSI values (*rbm-25* PSI – reference PSI; where ΔPSI<0 indicates decreased inclusion in *rbm-25* mutants; **Fig. 7B**). Alternative exons in the first three bins (>=100 nucleotides in length) demonstrate no significant change in PSI in *rbm-25* mutants. Beginning at bin 4 (exon size: 52-99bp) and becoming more pronounced in bins 5 and 6 (exon size: 30-51bp, and small microexons: <=27bp), there is a striking and significant decrease in exon inclusion in *rbm-25* mutants relative to the reference control samples (bin 4 (52≤exons ≤99) compared to bin 6 (microexons), pairwise Wilcoxon Rank-sum test, p = 1.7 x 10^-7^).

Next, we compared the size distribution of *rbm-25* regulated cassette exons with those of other known microexon regulators and other more broadly acting splicing factors (**Fig. 7C**). We identified all the cassette exons that were significantly differentially regulated (|ΔPSI| >=0.2) in five RBP mutant samples compared to their corresponding reference controls across several datasets. From *C. elegans*: *rbm-25*, *prp-40*, *mbl-1* and *exc-7*; from mouse, SRRM4/nSR100 was analyzed. PRP-40 is a known regulator of microexon splicing in worms (Choudhary et al., 2021). MBL-1 and EXC-7 are RNA-binding proteins expressed in the *C. elegans* nervous system that associate with intronic sequences downstream of an *unc-13* microexon and may facilitate recruitment of PRP-40 to promote its inclusion (Choudhary et al., 2025; Napier-Jameson et al., 2024). Lastly, SRRM4/nSR100 is a well-established regulator of microexon splicing in vertebrates (Irimia et al., 2014a; Quesnel-Vallières et al., 2015). When comparing the exon size distributions upon loss of each RBP (**Fig. 7C**), exons regulated by SRRM4, *rbm-25*, and *prp-40* exhibited a similar size distribution biased toward shorter lengths, with median sizes of 30 bp (IQR: 18–67), 36 bp (IQR: 24–52), and 39 bp (IQR: 26–72), respectively. Comparison of RBM-25-regulated exon sizes to those regulated by SRRM4 and PRP-40 revealed no significant differences in size distributions (Wilcoxon rank-sum test with Bonferroni correction: p = 1.0 and p = 0.7, respectively). In contrast, exons differentially spliced upon loss of *mbl-1* or *exc-7* had broader size distributions including larger exons (77 bp (IQR: 45-132) and 81 bp (IQR: 48-157), respectively), differing significantly from the *rbm-25* mutant profile (Wilcoxon rank-sum test with Bonferroni correction: p = 9.6e-05 and p = 8.2e-05, respectively). Collectively, these results indicate that microexons are generally more dependent on RBM-25 activity relative to longer alternative exons.

In yeast, Snu71 and Prp40 (RBM-25 and PRP-40 orthologs, respectively) are core components of the U1 snRNP, forming a stable subcomplex with Luc7 (Ester & Uetz, 2008; Gottschalk et al., 1998). In humans, RBM25 similarly copurifies with intact spliceosomes and U1 snRNP components (Fortes et al., 2007; Zhou et al., 2008), and though non-essential, exerts widespread influence on gene expression. Consistent with our results, *prp-40* has been identified as a key regulator of microexon splicing in *C. elegans* (Choudhary et al., 2021), we also found that *prp-40* and *rbm-25* EMS-recovered mutants led to reduction of microexon splicing reporter inclusion (**Fig. 5**), suggesting that both have roles in enhancing the inclusion of this microexon.

Upon observing that both RBM-25 and PRP-40 preferentially regulate microexons, we next asked whether these RBPs co-regulate the same set of microexons or act on distinct subsets. To address this, we calculated ΔPSI values for all detected microexons in *rbm-25* and *prp-40* mutant RNA-seq datasets relative to their reference controls (**Fig. 7D**). We found that the majority (85/114) of the measurable microexons exhibit a striking similarity in reduced inclusion levels in both *rbm-25* and *prp-40* mutants; only a small subset of microexons show increased inclusion in these mutants. This overlap among regulated target mRNA supports the idea that RBM-25 and PRP-40 likely function in the same pathway to achieve coordinated regulation of microexon inclusion.

Given that RBM-25 and PRP-40 and are both components of the U1 snRNP in yeast and humans, we wanted to test if these proteins physically interact as part of the same complex in *C. elegans*. We employed a co-IP (co-immunoprecipitation) strategy and collected lysates expressing an endogenously-tagged RBM-25::GFP fusion protein alone or co-expressing RBM-25::GFP and PRP-40::mScarlet fusions. Consistent with a role in the same protein complex, immunoprecipitation using α-RFP revealed that RBM-25::GFP co-IPs with PRP-40::mScarlet. Collectively our data demonstrates that RBM-25 and PRP-40 are both central regulators of microexon splicing in *C. elegans*, and likely act via their physical association with each other and the U1 snRNP to preferentially coordinate the inclusion of microexons.

## Discussion

During the course of normal development, changes in alternative splicing programs facilitate the development of tissues and the establishment of tissue identities. Uncovering the regulatory mechanisms governing splicing outcomes and demonstrating the functional consequences of individual splicing events as well as networks of co-regulated splicing events are major objectives within the field of alternative splicing. In order to address these questions, we undertook a large-scale exploration of alternative splicing in three tissues over the course of worm development from early embryo to adult, generating the most comprehensive metadata set of regulated splice variants across major tissue types in *C. elegans*. We delineate trends in developmental- and tissue-regulated alternative splicing, highlight potential RBPs coordinating similarly regulated splicing events and related processes and provide hypotheses for the investigation of individual splice variants.

### RNA regulons shaped by alternative splicing networks during C. elegans development

By probing patterns of gene expression (GE) and alternative splicing (AS) regulation, we reaffirm the roles of each in contributing to tissue identity during development, corroborating the idea that splicing is an additional layer of complexity specifying inter- and intra-tissue differences throughout development. Given our understanding of the complex crosstalk between AS and a number of other factors such as the chromatin landscape, transcriptional dynamics, non-coding RNAs, RNA editing and modifications (Luco & Misteli, 2011; Naftelberg et al., 2015; Shenasa & Hertel, 2019; Tellier et al., 2020), the study of AS regulation continues to reveal intriguing insights into tissue development and function. This dataset provides a high-resolution collection of splicing events that can leveraged to probe the interplay between splicing and other areas of gene regulation.

Examining the subset of genes that undergo developmentally-regulated splicing, we find that most are also doubly regulated in a tissue-specific manner, indicating that isoforms contributing to developmental identity also specify tissue-specific differences in splicing. Given the GO terms highlighted for each tissue, it will be interesting to explore how specific splice isoforms modulate protein activity and contribute both to specific functions within a developmental stage and within a specific tissue. In fact, GO terms in neurons such as *synaptic vesicle budding* (Blue et al., 2018), *axon extension* (Zheng, 2020) and *regulation of calcium ion transport* (Giudice et al., 2014) have each been investigated in the framework of the mammalian nervous system as well as other model organisms.

Groups of functionally related transcripts and/or transcripts that drive transitions during tissue developments have been demonstrated to be coordinated by shared regulatory mechanisms, often referred to as ‘RNA regulons’ (Baralle & Giudice, 2017; Keene, 2007). These regulons often involve single RBPs or a set of RBPs acting in a combinatorial manner to direct splicing programs (Kalsotra et al., 2008; Weyn-Vanhentenryck et al., 2018). We explored this concept by clustering tissue-regulated splicing events by the developmental stage(s) where they exhibit the greatest differential splicing, and therefore, likely instruct stage-specific tissue differences. Several of these exon junction usage trajectories contain genes that are interconnected in protein-protein interaction networks, aligning with the evidence that proteins regulated by tissue-specific splicing often demonstrate more extensive interaction networks (Buljan et al., 2012; Ellis et al., 2012; Kjer-Hansen & Weatheritt, 2023; Roth et al., 2023; Ule & Blencowe, 2019). Additionally, RBP motifs enriched in these splicing events provide a list of splicing regulators that are potentially regulating networks of interconnected genes. Furthermore, the enrichment of RNA-binding protein (RBP) motifs within these splicing events yields a catalog of potential splicing regulators that may be coordinating networks of interconnected genes. Further investigation into the regulatory role of these RBPs in shaping the splicing patterns of these events promises to shed light on their functions within specific biological processes.

### An expanded repertoire of microexons with distinct regulatory patterns

In this study, we discovered 176 microexons in the worm genome, over 40% of which were previously non-annotated. Although well-characterized in mammals, their prevalence, regulation, and biological relevance in *C. elegans* have not been well defined. Here, we integrate a computational framework for microexon detection with genetic screening and molecular approaches to expand the catalog of microexons in *C. elegans* and explore their regulation. Using these methods, we have also identified RBM-25, which, along with its binding partner PRP-40, acts as a critical splicing factor that enhances microexon inclusion.

Given the diversity of tissue types and developmental stages in this transcriptomic dataset, it is not surprising to find that the vast majority of microexons demonstrate some variation in inclusion across different samples. However, it is worth noting that the associated enriched GO terms of microexon containing genes in *C. elegans* were overwhelmingly related to neuromuscular development and phsyiology. This result is consistent with an earlier analysis of a smaller set of *C. elegans* microexons (Choudhary et al 2021). More broadly, this enrichment pattern mirrors the observations in mammals, zebrafish and fruit flies that most microexons switch inclusion states during neuronal maturation (Irimia et al., 2014b; Lopez-Blanch et al., 2025; Torres-Méndez et al., 2022), suggesting that microexon-based gene regulation has been co-opted in neural tissues across metazoa.

Individual microexon splicing events have been examined for their functional importance at various stages in neurodevelopment. Below we describe a subset of these findings, but other studies have been performed as well. For example, NOVA1/2-regulated inclusion of the 6b microexon in *Robo1/2* modulates downstream RAC1 and CDC42 signaling to regulate axon guidance (Johnson et al., 2019). In neuritogenesis, rescuing inclusion of a 6-nt microexon in *Unc13b* was sufficient to restore neurite growth in nSR100-deficient neurons (Quesnel-Vallières et al., 2015). Inclusion of E8a microexon in LSD1 (KDM1A) generates a neuronal isoform with altered transcriptional regulatory activity that promotes neurite outgrowth and maintains normal neuronal excitability (Rusconi et al., 2015). Microexons also regulate the function of several key structural proteins such as ankyrin-G, which has recently been shown to contain a highly conserved microexon E35a that is skipped in excitatory glutamatergic neurons but included in inhibitory GABAergic interneurons and fine-tunes Ca^2+^ signalling and neuronal excitability (Alam et al., 2026). The inclusion of a microexon in the formin gene, *Daam1*, extends the FH2 domain linker region, affecting a change in actin polymerization activity and influencing many downstream processes including dendritic spine formation, long-term potentiation and memory (Poliński et al., 2025). A 5-amino acid microexon included *Intersectin-1* reshapes its SH3A domain and changes its protein interactions to potentially regulated to regulate clathrin-mediated endocytosis of synaptic vesicles and synaptic transmission (Gerth et al., 2019).

In *C. elegans*, UNC-13, a highly conserved protein that facilitates synaptic vesicle release, contains a 9-nt microexon that demonstrates cell-type specific inclusion in the worm nervous system (Choudhary et al., 2025). Mediated by the RBPs, PRP-40, MBL-1 and EXC-7, this microexon is skipped in olfactory neurons, but included in motor neurons and appears to tune the strength of synaptic transmission according to the functional requirements of the cell-type. In this study, we have explored several microexon-containing genes with known roles in cellular architecture, axon guidance/neuronal identity (*madd-4*; **Fig. 5A**, (Pinan-Lucarré et al., 2014; Seetharaman et al., 2011)), ion channel-mediated signaling (*kvs-5*; **Supplemental Fig. 4**; (Hobert, 2013)) and neurodegeneration (*clp-1*; **Fig. 4E**, (Syntichaki et al., 2002)). We also highlighted the spatiotemporal regulation of microexon inclusion and the potential for these events to generate context-dependent protein functions. Future work will be required to determine how individual microexons modulate protein activity in specific cellular and developmental contexts.

An analysis of the GO terms associated with genes harboring microexons also revealed an enrichment for *mRNA splicing* related genes. Among the microexon-containing genes that belong to this GO term are *rbm-25* and *prp-39*, both components of the U1 snRNP and both containing putative, non-annotated microexons. The *rbm-25* microexon (22nt) falls immediately downstream of the N-terminal RRM domain within an intrinsically disordered region, while the *prp-39* microexon (5nt) falls within a TRP (tetratricopeptide repeat) domain, which typically modulates protein interactions. Inclusion of these micreoexons is predicted to result in downstream premature termination codons. Interestingly, in a forward genetics screen in *A. thaliana* searching for splicing regulators, mutations in *prp39a* (one of the two paralogs of yeast PRP39) led to splicing changes in its own transcripts and in *prp40b* transcripts (Kanno et al., 2017). Additionally, knockdown of PRP40A in mouse neuroblastoma cells increases the productive splicing of Luc7l, another U1 snRNP component, from approximately 30% under normal conditions to 80%, due to the increased skipping of a “poison exon” (Choudhary & Norris, 2024). Perhaps similar mechanisms of autoregulation are occurring in U1 snRNP components containing microexons in *C. elegans*. Splicing-dependent autoregulation has also been described for several other splicing factors, most notably the human SR protein family, in which each member contains a “poison exon” whose inclusion introduces an in-frame stop codon, targeting the transcript for nonsense-mediated decay (NMD) (Lareau et al., 2007). Examples of SR proteins regulating the inclusion of “poison exons” in other SR proteins has also been described (Leclair et al., 2020). Numerous core spliceosomal proteins are also autoregulated through alternative splicing coupled to nonsense-mediated decay, highlighting the importance of this mechanism as a means of fine-tuning the expression of spliceosome components (Saltzman et al., 2008).

In characterizing the fitness and behaviour of *rbm-25* mutants, we hypothesized a connection between microexon-containing genes that had an observable decrease in inclusion and the observed mutant phenotypes. For example, *rbm-25* mutants exhibited a 26% reduction in brood size, reduced number of progeny for the first three days and a trend towards less embryonic viability (**Fig. 6A**, **Supplemental Fig. 8A-B**). Numerous microexon-containing genes with reported activity in fertility, embryonic development, vulval development and egg laying were found to have reduced inclusion levels in mutants (ex. *clp-1* and *unc-13* mentioned previously). Similarly, mutant worms exhibited changes in locomotory patterns relative to wild-type (**Fig. 6C-D**). Although the molecular basis of these behavioural differences remains unclear, our data reveal RBM-25-dependent regulation of microexons in several genes encoding neuropeptides/receptors, and ion channels, providing potential candidates through which altered splicing may influence locomotion (ex. *bas-1* (Hare & Loer, 2004), *tyra-2* (Donnelly et al., 2013), *pezo-1* (Fazyl et al., 2025)). Lastly, the aldicarb resistance seen in *rbm-25* mutants, coupled with a normal response to levamisole, likely indicates defects in presynaptic ACh release. Many microexon-containing genes directly regulate synaptic vesicle formation, release and trafficking including *unc-13*, *dpy-23* (Yook et al., 2001), *tom-1* (Dybbs et al., 2005) and *dyn-1* (Parker et al., 2007). Future studies will be important to define how individual microexons modulate protein function, uncover the regulatory mechanisms that coordinate their inclusion, and determine how their misregulation contributes to altered tissue and developmental phenotypes.

### A specialized role PRP-40/RBM-25 containing U1 snRNPs, assisted by tissue-restricted RBPs, in microexon regulation

In *S. cerevisiae*, the U1 snRNP is composed of 10 proteins including Prp39, Prp40 and Snu71/RBM25, all three of which are essential for viability (Gottschalk et al., 1998; Lockhart & Rymond, 1994). Mutations in Prp39 and Prp40 lead to defects in alternative splicing in yeast (Gottschalk et al., 1998; Kao & Siliciano, 1996; Lockhart & Rymond, 1994), while the Snu71 homolog, RBM25, has been implicated in alternative splicing in humans (Carlson et al., 2017; Zhou et al., 2008). U1 snRNP binds to the 5’ splice site as the first critical step in the formation of the spliceosome (Lerner et al., 1980; Wahl et al., 2009b) and depletion of U1 levels results in changes to splice site recognition and exon inclusion (Dvinge et al., 2019). We postulate that a subset of U1 snRNPs containing PRP-40 and RBM-25 have been co-opted in *C. elegans* to preferentially support microexon inclusion. However, this mechanism of action is likely also conserved in vertebrates, as depletion of PRPF40A in mouse neuroblastoma cells can also lead to reduction of neural microexon inclusion (Choudhary & Norris, 2025) Consistent with the concept of specialized U1 snRNPs, a recent study has found that U1 snRNPs containing different paralogs of LUC7 proteins (LUC7L, L2, and L3) can regulate distinct alternative splicing outcomes by interacting with different 5’ splice site sequences (Kenny et al., 2025). Interestingly, elevated levels of RBM25 or Luc7L3 are sufficient to switch the splicing patterns of SCN5A transcripts, encoding a cardiac voltage-gated sodium channel, from the full-length isoform in normal conditions to truncated variants upon heart failure, resulting in a drop in the main current for conduction in cardiomyocytes (Gao et al., 2011). Thus, further study will be required to explore how PRP-40 and RBM-25 cooperate with each other, and perhaps with other U1 snRNP-associated factors, to preferentially regulate microexons over larger alternative exons.

Our microscopy data (**Fig. 5**) indicates that RBM-25::GFP is a widely expressed, nuclear protein. This raises the interesting question of how certain microexons are included in tissue-biased patterns. Our motif analysis also showed an enrichment in binding sites for the tissue-biased RBPs EXC-7, ASD-1 and FOX-1 in the introns downstream of microexons. Previous studies have established that the binding of RBPs to the downstream intron can promote the inclusion of conventionally sized alternative exons by stabilizing the U1 snRNP (Witten & Ule, 2011) and perhaps a similar mechanism is applicable for microexon splicing. It has been shown that *prp-40* genetically and physically interacts with *exc-7* and *mbl-1* to regulate microexon splicing (Choudhary et al., 2021, 2025). Additionally, *asd-1* and *fox-1*, homologs of the RBFOX family of splicing factors in humans that regulate microexon splicing in the nervous system (Y. I. Li et al., 2015b), were also shown to regulate subsets of PRP-40-mediated microexon splicing events (Choudhary et al., 2021). We also saw an enrichment in the motif for *sap-49*, which is involved in the binding of the U2 snRNP to the branch site (Champion-Arnaud & Reed, 1994). However, UG-rich sequences are also recognized by the CELF family of RBPs, UNC-75 and ETR-1, which have restricted expression patterns in neurons and muscle cells, respectively, and could serve as regulators (Kuroyanagi, Watanabe, Suzuki, et al., 2013; Milne & Hodgkin, 1999). Consistent with this possibility, we recently identified that UNC-75 was required for the proper neuronal inclusion on an 18 nucleotide microexon in transcripts from the *dpy-23* gene locus (Bhatnagar et al., 2026). We thus propose a model where RBM-25 and PRP-40 containing U1 snRNPs are required, but not sufficient, to drive microexon inclusion, assisted by the recruitment of one of several auxiliary sequence-specific RBPs binding in the downstream intron. How a distributed group of RBPs can each facilitate the recruitment of PRP-40 and RBM-25 will be an interesting problem to understand.

Human RBM25 was determined to bind to poly-G sequences (Carlson et al., 2017; Sénéchal et al., 2023; Van Nostrand et al., 2020), such as the Bcl-x exonic sequence motif 5’-CGGGCA-3’ (Zhou et al., 2008). However, our motif analysis did not detect any enrichment of G-rich sequences in and surrounding RBM-25 regulated microexons. Characterization of RBM25/RED120 in human cultured cells has demonstrated that it is a regulator of alternative splicing (Carlson et al., 2017). Additionally, tethering RBM25 to a deactivated Cas13 variant (dCasRx) was recently demonstrated to excel as an RNA-directed activator of exon inclusion, significantly increasing the inclusion levels of >90% of target alternative exons, including microexons (J. D. Li et al., 2024). These results suggest that RBM-25 has conserved activity in stimulating exon inclusion when positioned in the vicinity of an alternative exon.

The major regulator of microexon inclusion in mammals, SRRM4/nSR100, and its paralog SRRM3, binds to UGC-rich sequences in upstream introns flanking microexons, and contains a specialized eMIC domain that is necessary and sufficient to activate microexon inclusion (Gonatopoulos-Pournatzis et al., 2018; Torres-Méndez et al., 2019). SRRM3/4 appears to facilitate the formation of early spliceosome complexes by promoting the recruitment of SF1 and U2AF to the branchpoint sequence, followed by U2 snRNP (Torres-Méndez et al., 2019). It will be interesting to explore how RBM-25 achieves microexon inclusion mediated instead by U1 snRNP at the 5’ splice site. Given that nematode worms have likely lost an SRRM3/4 homolog in their evolutionary lineage, it will be interesting to understand whether RBM-25-mediated regulation represents a compensatory strategy evolved for microexon regulation, or whether it is a more ancient mechanism. The fact that loss of PRP40A activity leads to exon skipping of microexons in a mouse neuroblastoma line suggests that some mammalian microexons require U1 snRNP-associated components that we have identified in this study.

### Dynamic nuclear foci as sites coordinating microexon and alternative splicing?

Our microscopy with endogenously tagged RBM-25::GFP indicates that RBM-25 localizes to dynamic nuclear foci (**Fig. 5F**). This subcellular distribution is reminiscent of the localization of RBM25 to nuclear speckles in humans (Fortes et al., 2007; Zhou et al., 2008). Nuclear speckles are hubs for numerous RNA processing steps, particularly splicing, modification, packaging and export (Galganski et al., 2017). Human RBM25 localization to nuclear speckles was demonstrated to be dependent on the central R/E/D-rich region (Zhou et al., 2008). Recent results suggest that specific gene loci are recruited proximal to nuclear speckles, and this chromosomal clustering can impact splicing efficiency. Interestingly, in *C. elegans*, localization to dynamic nuclear foci has also been observed for UNC-75/CELF (Loria et al., 2003; Pilaka-Akella et al., 2025). However, it has been suggested that nuclear speckles, as they appear in specific vertebrate lineages, are not likely to form in *C. elegans* (Małszycki et al., 2026; Pham et al., 2021). Nevertheless, it will be interesting to determine whether other tissue-biased splicing regulators also display localization patterns in these dynamic nuclear sub-domains. More broadly, future work aimed at understanding the relationship between sites of transcription, microexon splicing dynamics, and their correlation with these spatial compartments will shed further light on these observations.

## Methods

### Obtaining Gene Expression and Alternative splicing data

RNA-seq datasets were obtained for intestine, muscle and neurons in embryonic, L4 and Adult stages of *C. elegans* development; NCBI BioProjects PRJNA477006 (Warner et al., 2019), PRJNA416495 (Gracida et al., 2017), and PRJNA400796 (Kaletsky et al., 2018). Embryonic data was collected every 90 minutes for five timepoints starting at mid-gastrulation and continuing until the embryo progressed to the threefold stage marked by the initiation of terminal cell differentiation (Warner et al., 2019). RNA for embryo and adult tissue samples was collected using Fluorescence-activated cell sorting (FACS)-sorting of labelled cells, while RNA for L4 stage animals was collected using the translating ribosome affinity purification (TRAP) method (Gracida & Calarco, 2017). STAR was used to align RNA-seq data using the attribute (--twopassMode), which allows increased sensitivity of mapping to novel junctions. Majiq (Vaquero-Garcia et al., 2016) was run using the resulting .bam/.bai files with the standard parameters and the attribute (-- disable-denovo-ir). Further downstream analysis was performed using Majiq to obtain lists of alternative splicing (PSI (percent spliced in)) and differential splicing (dPSI) events. These events were subsequently filtered for read count support and reproducibility as described in (Koterniak et al., 2020), to generate lists of high confidence splicing events. In general, splicing events that are considered alternatively spliced have PSI values >=0.05 and <=0.95, and events that are considered significantly differentially spliced between two conditions have a minimum dPSI value of ∼|0.20|. In parallel, HTSeq (Putri et al., 2022) was used to obtain raw gene counts using the same .bam files; the resulting counts were normalized and log2 transformed using DESeq2 (Love et al., 2014) for downstream analyses. Genes that have >=50 supporting reads in one or more samples were considered as “expressed”.

### Principal Component Analyses (PCA) and Correlation plots

PCA analyses were performed on gene expression (z-score standardized, log2 transformed normalized gene counts) or alternative splicing (PSI) data obtained as described above. Based on the clustering of the replicates in diagnostic PCA analyses containing all the available data, the following replicates were excluded from downstream analyses for both gene expression and splicing due to divergence from the other replicates: embryo muscle replicates 1 (SRR7443653, SRR7443652, SRR7443651, SRR7443650, SRR7443649) and 2 (SRR7443648, SRR7443647, SRR7443646, SRR7443656, SRR7443655) at all five timepoints, and adult muscle replicate 3 (SRR7443988). Pearson correlation matrices were performed on the same two datasets. Pearson correlation graphs displaying the relationship between later developmental timepoints relative to the early embryonic T0 timepoint excluded the L4 stage data. All plots were generated in R.

### Comparison of differential splicing events between developmental stages and tissue types

Developmentally-regulated (D-R) events refer to splicing events exhibiting significantly differential usage patterns between two developmental stages within the same tissue. Tissue-regulated (T-R) events involve splicing events with differential usage patterns between two tissues at the same developmental stage. Significantly differential D-R splicing events (dPSI=∼|0.2|) were obtained for two pairwise comparisons for each tissue: 1) early vs. late embryo, 2) late embryo vs. adult. Each D-R splicing event was represented by a single junction that demonstrated the largest difference in PSI between conditions. Corresponding dPSI values of these junctions was obtained from T-R data. A heatmap was generated sorted by dPSI values of the D-R events. The colour scale of the heatmap was arranged such that events with a higher LSV (Local Splice Variation) usage in the earlier developmental stage are shown in green and those with a higher LSV usage in the later developmental stage are shown in purple. Conversely, for T-R events, purple represents high LSV usage in the tissue of interest and green represents high LSV usage in the other two tissues.

D-R events were grouped for each tissue and GO (Gene Ontology) term enrichment analyses of all the genes with significantly differential splicing was performed using ClueGO (Bindea et al., 2009).

In the Venn diagrams, unique genes that are significantly D-R in each pairwise developmental stage comparison for the tissue of interest were contrasted to the list of unique genes that are significantly T-R (pairwise comparisons between the tissue of interest and the two other tissues).

### Tissue-regulated splicing over developmental time

For any given pairwise tissue comparison, significantly differential T-R splicing events were obtained for early embryonic, late embryonic, L4 and Adult stages. A single local splice variation (LSV) junction corresponding to the largest dPSI was selected for each splicing event. The dPSI values for this junction were obtained from the pairwise tissue comparisons at all the other timepoints, thereby creating a list splicing events that are significantly T-R in at least one timepoint. These lists were filtered for splicing events that have reportable (non-NA) values for at least three of the four developmental stages. dPSI values were converted to absolute values and are referred to as “differences in isoform usage” as any given value does not necessarily correspond to a junction representing the “exon included” isoform. Data was clustered using Cluster 3.0 (kmeans, where k=8 or 9) and plotted in R.

Splice isoforms were sorted into ten select trajectories over developmental time and plotted in R, where each red line represents one splicing event, the blue line is the regression line, and the grey shaded areas are the 95% confidence intervals. For each trajectory, GO term over-representation analyses were performed on the list of genes present using WebGestalt (Elizarraras et al., 2024), and the highest scoring terms in biological processes, cellular components and molecular functions were reported. Protein-protein interaction network analysis was also performed in STRING (Szklarczyk et al., 2023) and filtered in Cytoscape (Shannon et al., 2003) for genes with at least one interaction. Motif enrichment analysis was performed using MEME-SEA (Bailey & Grant, 2021) to identify known RBP motifs from the CISBP-RNA database (Ray et al., 2013) within the transcript sequences containing T-R splicing events.

### Culturing of C. elegans strains and strains used

All phenotyping assays and growth of animals were performed with animals maintained for several generations at 21C and grown on 35 mm nematode growth medium (NGM) agar plates seeded with *Escherichia coli* OP50-1 bacteria under standard laboratory maintenance conditions for *C. elegans* (Brenner, 1974) unless otherwise noted. Key strains used in this study were N2 (wild-type), *smg-1(r861) I; csbIs36 [rgef-1_p_::madd-4 exon 6 two-colour reporter::tbb-2 3’UTR], smg-1(r861) I; rbm-25(csb87) V; csbIs36, smg-1 (r861) I; rbm-25(csb88) V; csbIs36, smg-1(r861) I; rbm-25(csb89) V; csbIs36.* All additional strains used in this study are available upon request.

### Construction of two-colour splicing reporters, microinjection and microscopy

Two-colour reporters were generated, injected, stably integrated and imaged using confocal microscopy as described in (Koterniak et al., 2020). The following promoters were used to drive expression in the tissues of interest, at the required developmental stages: *cnd-1* (embryonic neurons), *rgef-1* (L4 and Adult neurons), *myo-3* (L4 and adult body-wall muscle). All protein domain identification to map alternative exons onto encoded protein sequences was performed with InterProScan (Jones et al., 2014).

### Genome-wide microexon identification

RNA-seq datasets were obtained for embryonic, L4 and Adult stages of *C. elegans* development; NCBI BioProjects PRJNA477006, PRJNA416495, PRJNA400796. The embryo data contains transcriptomes from seven cell/tissue types, L4 data for five cell/tissue types and adult data for four tissue types. STAR and OLego mapping tools were implemented using the parameters outlined in (H. Yu et al., 2022). The reference genome fasta file and the GFF3 genome annotation file (genome assembly version: WBcel235, and gene annotation version: WBPS17) were used to generate STAR genome indexes (STAR runMode genomeGenerate). Subsequently, 1^st^ pass mapping of RNA-seq fastq files was performed with STAR (--alignIntronMin 20 -- alignIntronMax 20000 --outSAMtype None --outSJfilterReads Unique -- outSJfilterCountUniqueMin 10 3 3 3 --outSJfilterCountTotalMin 10 3 3 3). In parallel, OLego was used to build a genome index file using the same reference genome fasta file (-a bwtsw), then RNA-seq data was mapped without the genome annotation (-e 3 -I 20000 --max-multi 5). Sorted .sam files were converted to .bed files then to .junc files with the sam2bed.pl and bed2junc.pl scripts respectively, found in the OLego package. STAR SJ.out.tab files and OLego .junc files were each merged into a single file and used to regenerate the genome in STAR (--runMode genomeGenerate --genomeFastaFiles ref_genome.fa --genomeSAindexNbases 10 -- sjdbFileChrStartEnd star.SJ olego.SJ --sjdbOverhang 100 --limitSjdbInsertNsj 2000000). STAR 2^nd^ pass mapping was performed (--alignIntronMin 20 --alignIntronMax 20000 --outSAMtype BAM SortedByCoordinate --outSAMstrandField intronMotif --alignSJoverhangMin 20). StringTie (Pertea et al., 2015) was used for transcript assembly (.bam files) of all samples (stringtie -j 5 -c 5 -g 10 -G annotation_file.gff3) and the resulting individual .gtf files were merged to create a new annotation file. All samples were analyzed with Majiq (Vaquero-Garcia et al., 2016) in two separate runs: 1) using the standard annotation file and 2) using the StringTie merged annotation file. A list of all microexons <=27 nt were also extracted from the genome annotation file. Together, these three sets of microexons were filtered using the following parameters: 1) all first exons were removed. 2) all microexons that coincide with existing longer exons were removed. 3) all microexons were required to have junction support at both 5’ and 3’ ends, and at least two samples in the dataset were required to have >=10 reads. This resulted in the broad list of microexons, which was then further filtered for conservation of the dinucleotide donor and acceptor splice sites as well as the microexon proper (threshold for average PhyloP scores were >=1) to obtain a stringent microexon list. The same pipeline was used to analyze RNA-seq data for *rbm-25* mutants (this study)*, prp-40* (PRJNA684142), *exc-7* and *mbl-1* mutants (PRJNA386174).

### RT-PCRs and long-read nanopore sequencing of microexons

Total RNA was extracted from worms using TRIzol (Thermo Fisher Scientific) and chloroform followed by ethanol precipitation and RNA clean up/DNase treatment with the RNA Clean and Concentrator kit (Zymo Research Corp.). RNA concentration was quantified using a spectrophotometer and diluted to 10ng/ul. RNA was extracted from Bristol N2, *madd-4 two colour reporter; smg-1*, and *rbm-25; madd-4 two colour reporter; smg-1* strains. RT-PCR assays were performed with the OneStep RT-PCR kit (Qiagen) with 20ng of RNA per reaction as recommended by the manufacturer.

A set of 12, non-annotated microexons were selected from the stringent microexon list described above. Three primers were generated for each microexon: 1) forward primer annealing to the upstream exon (FwE1), 2) forward primer annealing to the putative microexon (FwMx), 3) reverse primer annealing to the downstream exon (RvE3). We validated the presence of microexon-containing transcripts using FwMx and RvE3 primer pairs. We validated the integrity of the RNA samples using the FwE1 and RvE3 primer pairs. We tested for DNA contamination of RNA samples by having a no-RT control (using the FwMx and RvE3 primer pairs), in which the first incubation of samples at 50°C for the reverse transcription step was skipped.

We performed long-read Nanopore sequencing (Plasmidsaurus) of unannotated microexons and high-confidence annotated microexons (located in the genes *cpna-2*, *dnc-1*, *noca-2*, *unc-104*, *lgc-40*, *kvs-5*, *exp-1*). Primers amplifying sequences between the upstream and downstream exons were used to perform RT-PCRs and these samples were purified and submitted for sequencing. For a list of all primers used in this study, please see **Supplemental Table S6**.

### EMS mutagenesis screen and whole genome sequencing

An ethyl methanesulfonate (EMS) mutagenesis screen was performed on the *madd-4 two colour reporter; smg-1* strain. Synchronized L4 larval worms were washed with M9 buffer and exposed to EMS at a final concentration of 0.05M in M9 buffer for 4 hours at 21°C with rotation. After washing worms twice with M9 buffer, mutagenized P_0_ worms were placed on OP50-containing plates and rested for ∼20 minutes. Twenty plates of 3-4 P_0_ worms each were grown until the F_1_ generation was gravid. F_1_ worms were washed with a 20μm mesh and diluted with NGM for a final concentration of 1 worm/200ul NGM. These worms were then singled into 20 x 96 well plates with added OP50, sealed with breathable sealing film and grown with agitation at room temperature until ready to screen when sufficient F_2_ worms were visible. F_2_ containing wells positive for a change in reporter phenotype were transferred to an OP50 plate and propagated by maintaining mutant heterozygous lines. EMS mutants were backcrossed to the parental *madd-4 two colour reporter; smg-1* strain 3-4 times.

In order to perform whole-genome sequencing to identify causal mutations, twenty homozygous mutant worms were picked into M9 buffer, allowed to rest for 30 minutes to remove residual OP50, pelleted and plated on unseeded plates for another 30 minutes. 15 worms were then picked into 10μl of PBS, placed at -80°C for 30 min. Genomic DNA extraction and amplification was performed with the REPLI-g Kit (Qiagen) as recommended by the manufacturer. Samples were purified using Quick-DNA Miniprep Plus kit (Zymo Research Corp.), biological fluids and cells protocol, and submitted for whole-genome sequencing (The Center for Applied Genomics (TCAG), SickKids Research Institute, Toronto, ON, Canada).

### rbm-25 CRISPR mutagenesis

CRISPR mutagenesis to generate the *rbm-25 in/del* mutant was performed according to the protocol found in (Martin & Calarco, 2022a). Two gRNA were designed targeting the start codon of *rbm-25*, transcribed, purified and incubated with Cas9, then subsequently injected into ten *madd-4* two colour reporter expressing animals. Mutant animals were then screened for two-colour reporter splicing defects. Site-directed mutagenesis via CRISPR to regenerate the EMS point mutation at the 3’ splice site flanking *rbm-25* exon 6 was performed using a single-stranded oligodeoxyribonucleotide (ssODN) as a repair template as described in (Martin & Calarco, 2022). The ssODN contains 35nt of homologous sequences on either side of a mutated restriction enzyme site used for genotyping and a mutated PAM sequence. This ssODN was co-injected with Cas9 and a gRNA targeting the same region. All mutants were also Sanger sequenced to verify that mutations were as expected.

### Confocal Microscopy

All microscopy was performed using a Leica SP8 laser scanning confocal. Two-color reporter imaging was performed with a 40x oil immersion objective, and RBM-25::GFP fluorescence imaging was performed with a 63x oil immersion objective. For imaging RBM-25::GFP nuclear foci, we applied the Lightning software for increased resolution of images collected over a time series of several minutes.

### Pharmacological and behavioral assays

To perform the aldicarb assay, a 100 mM stock solution was prepared in 70% ethanol and then added to NGM agar media to achieve a final concentration of 1 mM aldicarb. Plates were poured and seeded with OP50. 20-30 young adult worms from each genotype (blinded) were placed on each plate (in triplicates, performed over two separate dates). The number of paralyzed worms were scored every 30 minutes until all the worms in both phenotypes were paralyzed. State of paralysis was confirmed if the head and tail were each prodded three times with a worm pick but resulted in no forward or reverse movement. The protocol used was adapted from (Choudhary et al., 2025; Locke et al., 2008).

For levamisole paralysis assays, a levamisole 200 mM stock solution was prepared in water, and plates were poured with a final concentration of 0.5 mM levamisole. 20-30 young adult worms from each phenotype (blinded) were placed on each plate (in triplicates) and scored for paralysis every 30 minutes. Count data for both aldicarb and levamisole assays can be found in **Supplemental Table S5**.

### Image acquisition of motor behaviour on solid agar

All recordings for the motor behaviour of animals on solid agar were performed using a custom-built imaging setup in an unlit room and covered with a dark sheet. The imagine setup consisted of a glass plate suspended on four adjustable legs that were approximately 20 cm above a red LED strip light source. The trial plate of freely behaving animals was placed on top of the glass plate, such that the red LED strip below would provide evenly distributed darkfield illumination imaged by a USB camera positioned directly above the plate, capturing stop motion images at 4 frames per second (FPS). The camera used for recording locomotion data was the Mightex USB3.0 with the Aptina 5 Megapixel CMOS image sensor in monochrome. This camera was attached a Computar C-mount lens with a fixed focal length of 8 mm and an aperture range of F1.4 – F1.6.

L4 animals of each strain were selected 24 hours before assay, allowing them to mature to the young adult stage for behavioural assessment. Immediately prior to the assay itself, 2-3 animals at a time were aspirated into a 5 µL droplet of M9 buffer using a micropipettor with a plastic tip, transferring them within the droplet from 35 mm NGM plates seeded with OP50-1 to an unseeded intermediary 60 mm plate of NGM agar. 5 animals per behaviour assay trial were transferred by this method to an intermediary plate in order to clean off bacterial residue from the growth plate, and the intermediary plate was reused between different trials. The animals were then allowed to thrash for 3 minutes until the agar had absorbed the small liquid droplet and the animals started crawling again, upon which they were gently transferred using an eyelash pick onto the trial plate, which was also unseeded NGM agar poured into a 60 mm Petri dish. The trial plate additionally had a 35 mm boundary of 1 M CuSO4 solution stamped into the centre using a 3D printed ring-shaped piece. Copper is an aversive signal to *C. elegans* and the liquid solution stamping constrains the animals to a circular arena of movement with a 35 mm diameter, while avoiding imaging artifacts caused by physical boundary objects. When all 5 worms were transferred into the centre of the 35 mm CuSO4 ring on the trial plate, the trial plate was placed on top of the glass and images at 4 FPS were recorded for 10 minutes, resulting in a total of 2400 frames per trial. This process was repeated for as many trials as necessary to achieve the appropriate number of animals.

### Image acquisition of motor behaviour in liquid M9 buffer

Imaging of animals performing motor behaviour in a liquid medium by thrashing was recorded using 30 FPS smartphone cameras mounted to the eyepiece of a Zeiss Stemi 305 compact stereo microscope with brightfield illumination. Two young adult animals per trial were picked and transferred to a 35 mm trial plate of unseeded NGM agar according to the same method outlined in the animal transfer process for motor behaviour on solid agar. After 3 minutes of allowing the animals to acclimate and letting the small M9 buffer droplet to get absorbed into the agar, a larger droplet of 25 µL M9 buffer was pipetted directly on top of the two worms, and recording was started on the smartphone camera. The thrashing behaviour was recorded for 2 minutes uninterrupted, and this process was repeated for as many trials as necessary.

After image acquisition, these videos were uploaded to a computer and processed through the FFmpeg command-line tool to invert them (turning the animals from dark objects on a light background to light objects on a dark background), crop out the first minute of the video, and then convert the second minute of video into individual images at 30 FPS, resulting in 1800 frames per trial.

### Post-imaging analysis of motor behaviour in solid and liquid media

Raw behaviour data in the form of many frames with the animals as light objects on a dark background were processed using the FIMTrack v3 open-source software. After defining a region of interest (the area inside the CuSO4 ring for the solid medium behaviour assay and the extent of the M9 droplet for the liquid medium behaviour assay) and adjusting user-set parameters to match the FPS and minimum/maximum larvae area, the FIMTrack software automatically detects objects and tracks them across the provided images. The resulting tracks were manually curated to resolve colliding animals and other artifacts, with data compiled in the form of overall trajectory images and CSV tables containing the parameter value for each animal over each frame it was tracked. FIMTrack default parameters include centroid-based measurements useful for detecting average speed and distance travelled for animals behaving on a solid medium, as well as body curvature-based measurements useful for detecting coiling events and bending angle for animals behaving in a liquid medium. These data tables from FIMTrack were then processed further using a custom-written R script to calculate values for more *C. elegans*-specific parameters based on the default parameters measured by FIMTrack.

### Brood size assay

20 gravid adults per strain were picked onto a 35 mm NGM agar plate seeded with OP50-1 to perform a synchronized egg lay for 1 hour. Adults were picked off and the eggs remaining on the plate were incubated at 21C for two days to mature into L4 animals, upon which they were each singled onto their own growth plates. Single fertile animals were monitored and transferred onto a new growth plate daily, with the plate from the previous day kept and incubated for another 24 hours at 21C. These progeny-containing plates were then monitored the next day, with the total number of hatched larvae and unhatched progeny counted and recorded. This daily counting and transferring of animals continued until either the animal died or had not laid any progeny for two days in a row.

### Lifespan assay

Embryos from a plate full of gravid adults were transferred onto several fresh NGM agar plates with OP50-1. The embryos were incubated for two days at 21C until they matured into L4 animals, upon which they were picked onto separate growth plates at 5 animals per plate. Animals were monitored on a daily basis and lethargic animals were tested for survival by observing pharynx pumping, movement when the agar next to the head or tail of the animal was tapped, or movement when the head or tail of the animal was directly tapped by a platinum wire pick. Status of each animal was recorded in a spreadsheet, marking the daily count of animals that were alive, dead, or censored from the data due to burrowing into the agar or crawling up the side of the plate and desiccating.

### Co-immunoprecipitation and western blotting

Well grown plates (2-3 plates, 60 mm) containing mixed stage animals were washed 2-3 times with M9 buffer to minimize bacterial contamination. The collected animals were homogenized in RIPA buffer (rtak, Proteintech) supplemented with the protease inhibitor cocktail (Complete Mini EDTA free protease inhibitor, Roche) and 1 mM phenylmethylsulfonyl fluoride (PMSF) (Millipore Sigma). The homogenization was carried out at 4°C to prevent protein degradation. The supernatant was separated from the worm debris by centrifuging at 8000 x g in a tabletop refrigerated centrifuge. The supernatant was collected in a separate microfuge tube and incubated with RFP-Trap (ChromoTek RFP-Trap Agarose, Proteintech, Cat No. rtak) at 4 o C on a revolving rotator for 4-5 hours. Subsequent washing and elution steps were followed as described by the manufacturer. The final output was eluted in final volume of 80 ul. The final eluted samples were analyzed by using standard western blot protocol and probed for the following antigens using specific primary monoclonal antibodies: Mouse Anti-RFP monoclonal (6G6) (Proteintech) for mScarlet, rat Anti-GFP monoclonal (3H9) (Proteintech) for GFP. Secondary antibodies conjugated with horseradish peroxidase (HRP) are as follow: Anti-Mouse-HRP (CST), Anti-Rat-HRP (CST).

---

**Supplemental Figure S1:**
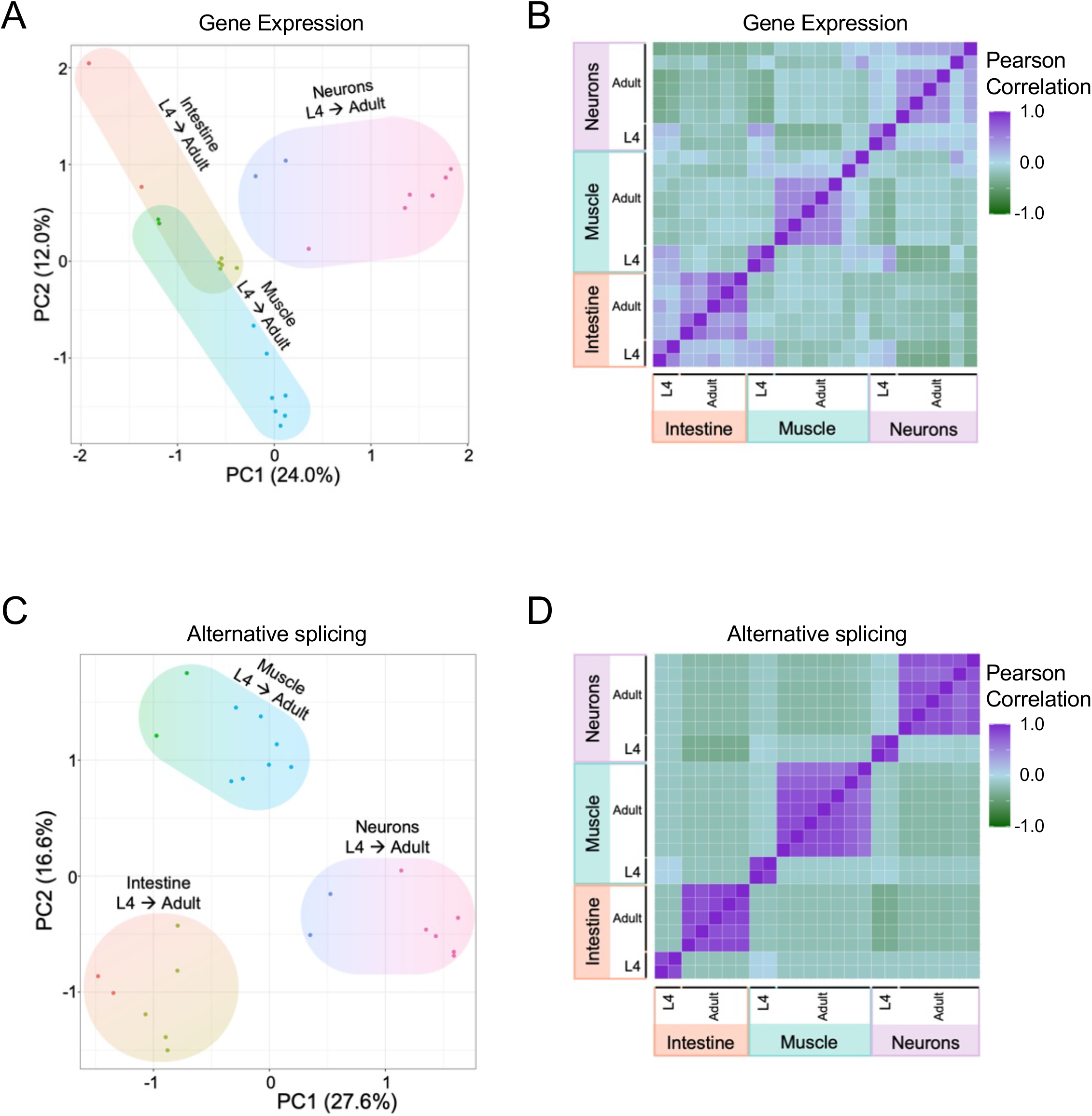
Correlation Tissue-specific gene expression and alternative splicing profiles across L4 and adult developmental time points. A) Principal Component Analysis (PCA) plot of normalized gene expression counts (log₂-transformed, standardized) at L4 (two replicates) and adult time points (5-7 replicates) in three tissues: intestine (orange–green), muscle (green–blue), and neurons (purple–pink). Arrows indicate the direction of L4 to adult progression. B) Matrix of pairwise Pearson correlations of the gene expression data described in A). Each cell shows the correlation between two samples, with purple indicating strong positive correlation and green, strong negative correlation. C) PCA plot of standardized percent spliced in (PSI) values of all the alternative splicing events detected in the data described in A) Arrows indicate the direction of L4 to adult progression. D) Matrix of pairwise Pearson correlations of the alternative splicing data described in C). Each cell shows the correlation between two samples, with purple indicating strong positive correlation and green, strong negative correlation.

**Supplemental Figure S2:**
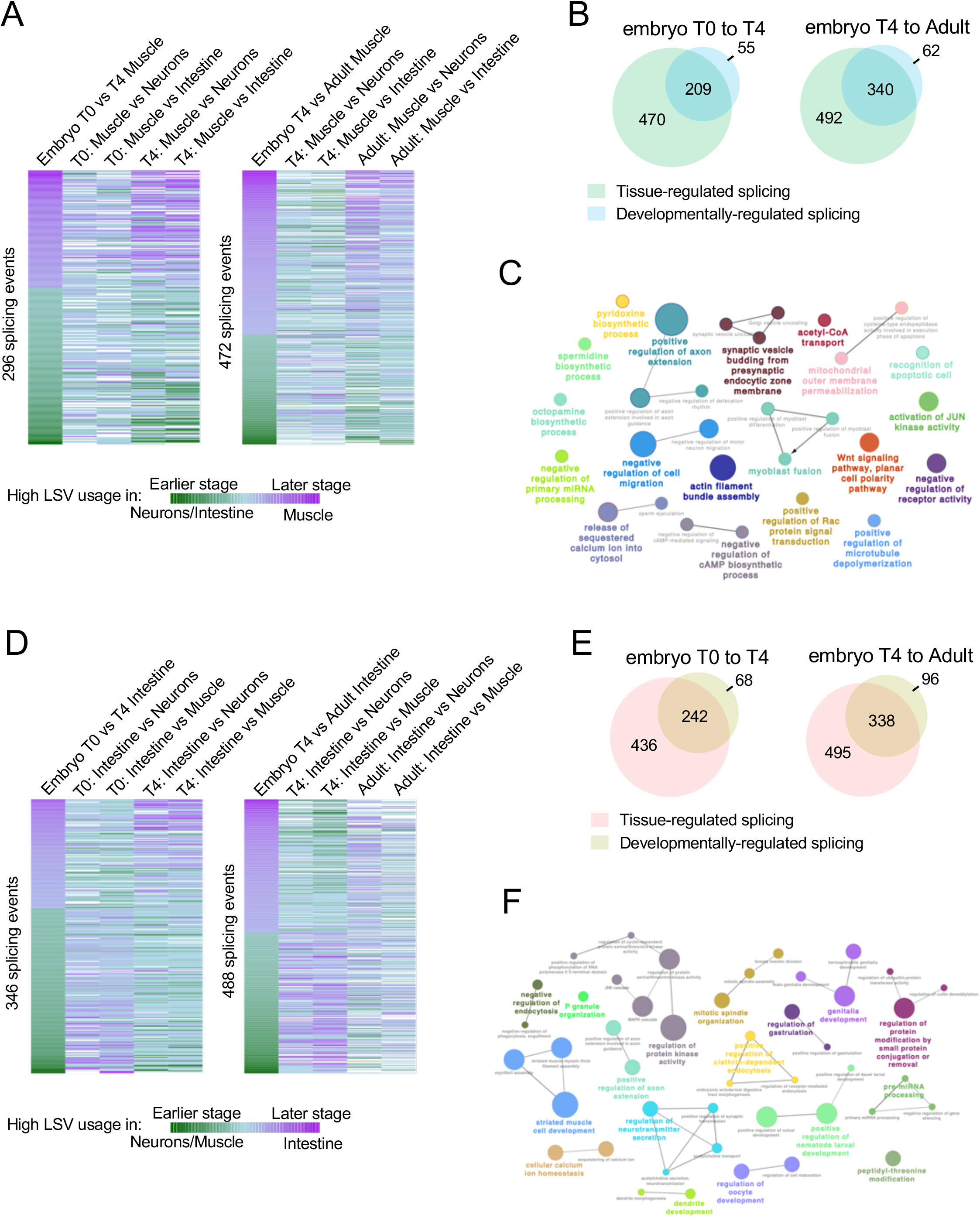
Interplay between developmental- and tissue-specific regulation of splicing programs. A) Left heatmap: 296 alternative splicing events (LSVs) with significant developmental regulation between early and late stages of embryonic muscle development (Embryo T0 vs. T4 muscle, left column), compared with LSV usage in pairwise tissue comparisons (muscle vs. neurons and muscle vs. intestine at the corresponding developmental stages, T0 and T4, remaining columns). For developmental regulation, green indicates higher LSV usage at T0 and purple at T4; for tissue comparisons, green indicates higher LSV usage in neurons/intestine and purple in muscle. Right heatmap: Comparing LSV usage in 472 developmentally-regulated splicing events between T4 and adult stage muscle with LSV usage values between tissues at the same stages. B) Venn diagrams showing the overlap between genes containing developmentally-regulated splicing events (blue; differences between two time points in muscle) and genes containing tissue-regulated splicing events (green; differences between muscle and and neurons/intestine at the same time points). Left diagram: embryo T0 and T4 comparison. Right diagram: embryo T4 and adult comparison. C) GO term enrichment analysis using ClueGO of developmentally-regulated genes in muscle (all developmental stages combined). D) Left heatmap: 348 alternative splicing events (LSVs) with significant developmental regulation between early and late stages of embryonic intestinal development (Embryo T0 vs. T4 muscle, left column), compared with LSV usage in pairwise tissue comparisons (intestine vs. neurons and intestine vs. muscle at the corresponding developmental stages, T0 and T4, remaining columns). For developmental regulation, green indicates higher LSV usage at T0 and purple at T4; for tissue comparisons, green indicates higher LSV usage in neurons/muscle and purple in intestine. Right heatmap: Comparing LSV usage in 488 developmentally-regulated splicing events between T4 and adult stage intestine with LSV usage values between tissues at the same stages. E) Venn diagrams showing the overlap between genes containing developmentally-regulated splicing events (blue; differences between two time points in intestine) and genes containing tissue-regulated splicing events (green; differences between intestine and and neurons/muscle at the same time points). Left diagram: embryo T0 and T4 comparison. Right diagram: embryo T4 and adult comparison. F) GO term enrichment analysis using ClueGO of developmentally-regulated genes in intestine (all developmental stages combined).

**Supplemental Figure S3:**
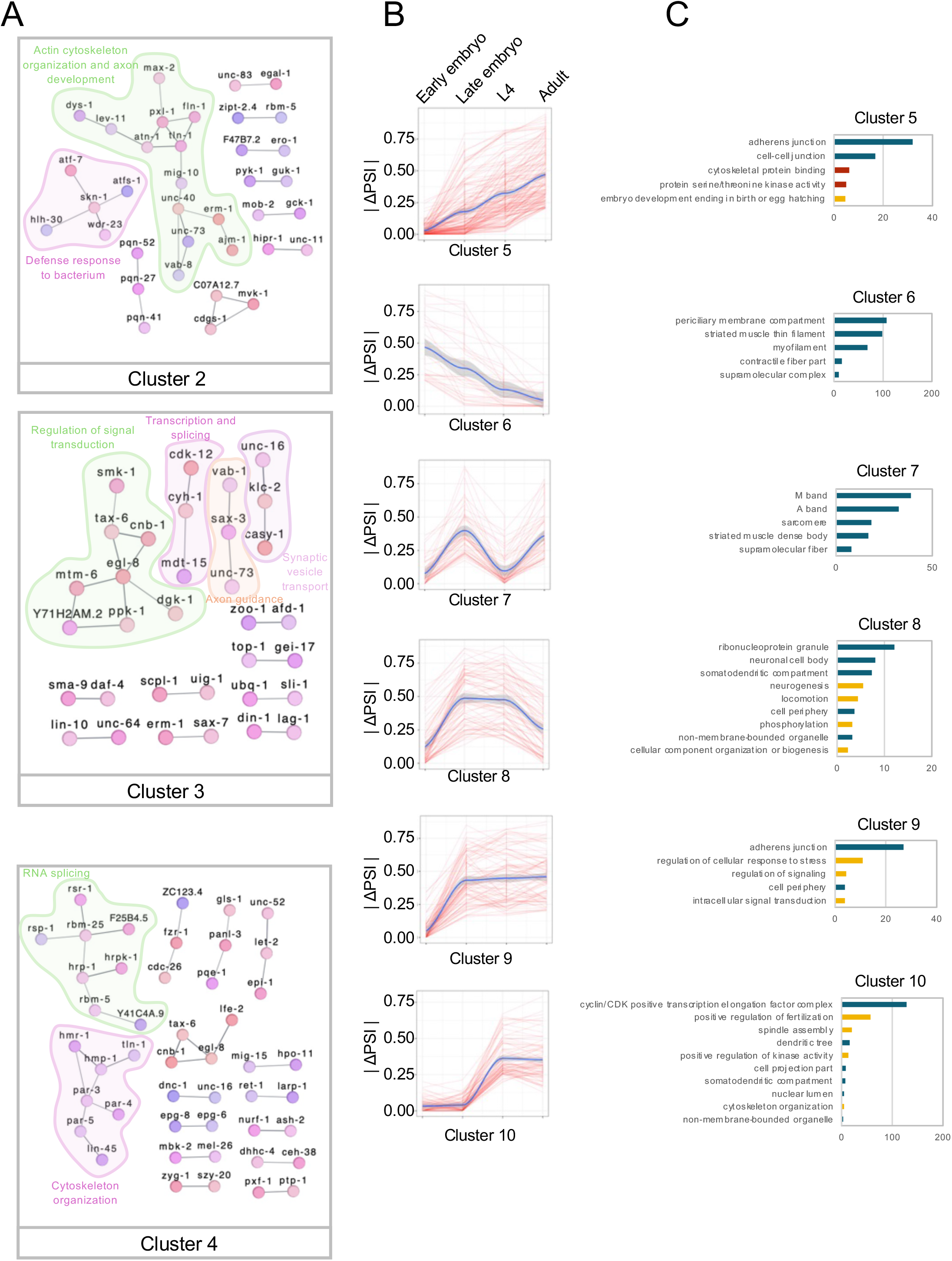
Network and splicing trajectories of tissue-specific splicing events across development. A) Interaction networks of three clusters of genes with tissue-specific splicing events that share a pattern of |Δ PSI| across development as seen in Figure 3E. Networks were obtained from STRING and show only genes with a minimum of one interaction. Applied high confidence interaction score = 0.700. B) Clusters 5-10 of splicing trajectories that group splicing events from all three heatmaps seen in Figure 3D that follow a set pattern of |Δ PSI| across development. Blue line is the regression line and the shaded grey area is the 95% confidence interval. Cluster 5 comprises all tissue-regulated splicing events that showed a linear increase in |Δ PSI| as development progresses from early embryo to adult. Opposingly, Cluster 6 events show linear decrease in |Δ PSI|; Cluster 7 events demonstrate a bimodal |Δ PSI| with peaks at the late embryo and adult stages; and Cluster 8 events show a |Δ PSI| peak at the late embryo and L4 stages; Cluster 9 events have high |Δ PSI| at the late embryo stage onwards and lastly, Cluster 10 events have high |Δ PSI| at the L4 stage onwards. C) GO term enrichment analysis on the events categorized in each cluster from B). Enrichment score bars are colored according to the three GO aspects: blue = Cellular Component, red = Molecular function, yellow = Biological Process.

**Supplemental Figure S4:**
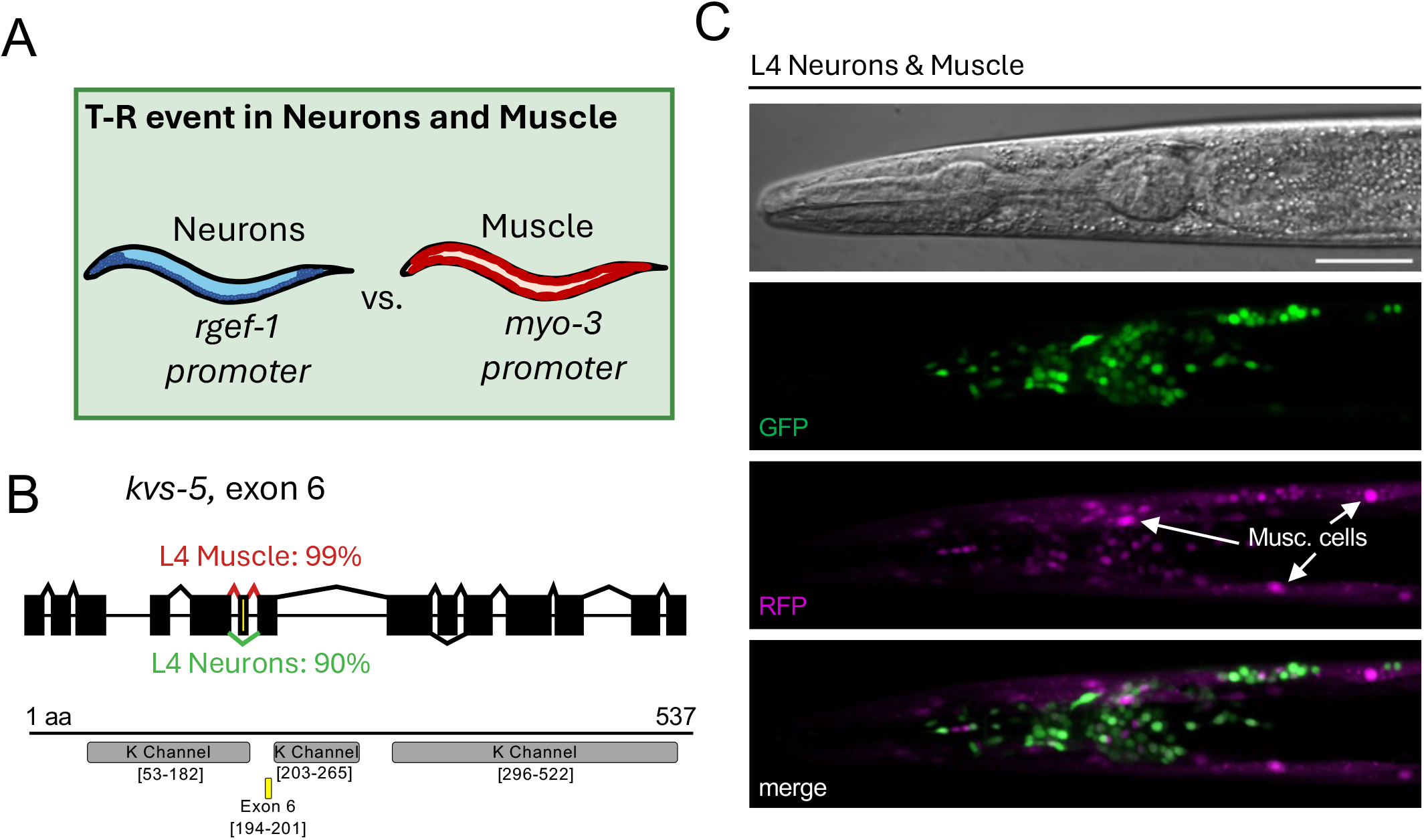
Two-colour reporter of a cassette exon demonstrating tissue-regulated splicing in neurons and muscle. A) Two-colour splicing reporters designed to explore splicing differences in neurons and muscle of L4 worms are expressed under the control of the driven by the *rgef-1* promoter for the former and *myo-3* promoter (body-wall muscle) for the latter. B) Map of the *kvs-5*, exon 6 cassette (microexon) splicing event predicted from RNA-seq data to be predominantly skipped (90%) in neurons and highly included (99%) in muscle. Diagram below indicates the position of conserved protein domains (grey) and the alternative exon (yellow) with bracketed ranges denoting the corresponding amino acid positions. C) Fluorescence microscopy of a worm co-expressing neuron-specific and muscle-specific splicing reporters for *kvs-5*, exon 6 at the L4 stage (scale bar = 50µm). RFP or GFP signals indicate the inclusion or skipping, respectively, of the alternative exon 6. Small, medially located puncta are neurons, while larger puncta are body-wall muscle cells a few of which are indicated by white arrows.

**Supplemental Figure S5:**
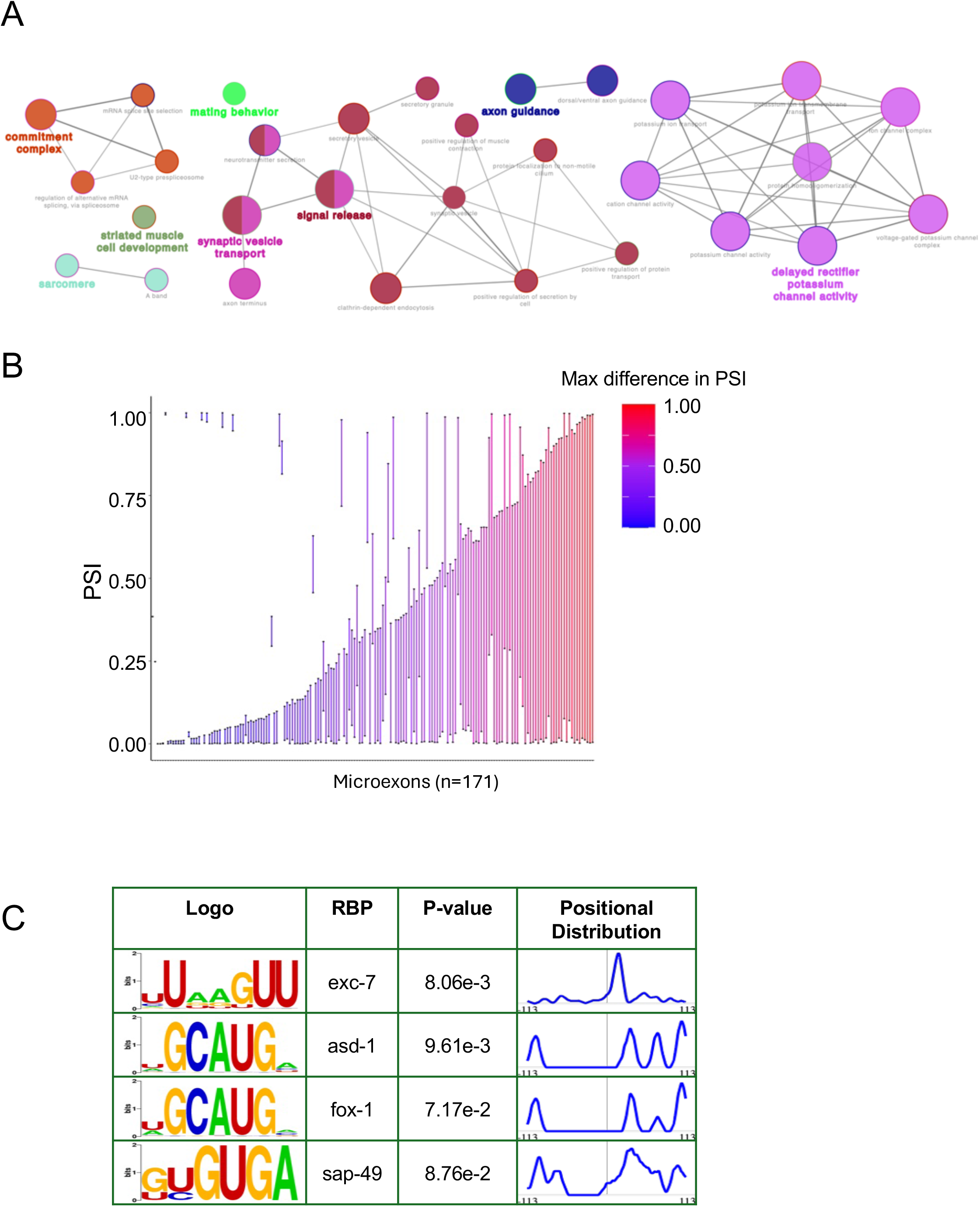
Features associsted with expanded repertoire of microexons in *C. elegans.* A) Enriched GO network based on the collection of genes containing microexons performed using ClueGO in Cytoscape. B) Graph showing the range (maximum–minimum) of PSI values for each microexon across RNA-seq samples, with colours indicating the magnitude of PSI change. C) MEME-SEA enrichment of known RBP motifs in microexon + flanking 100nt sequences.

**Supplemental Figure S6:**
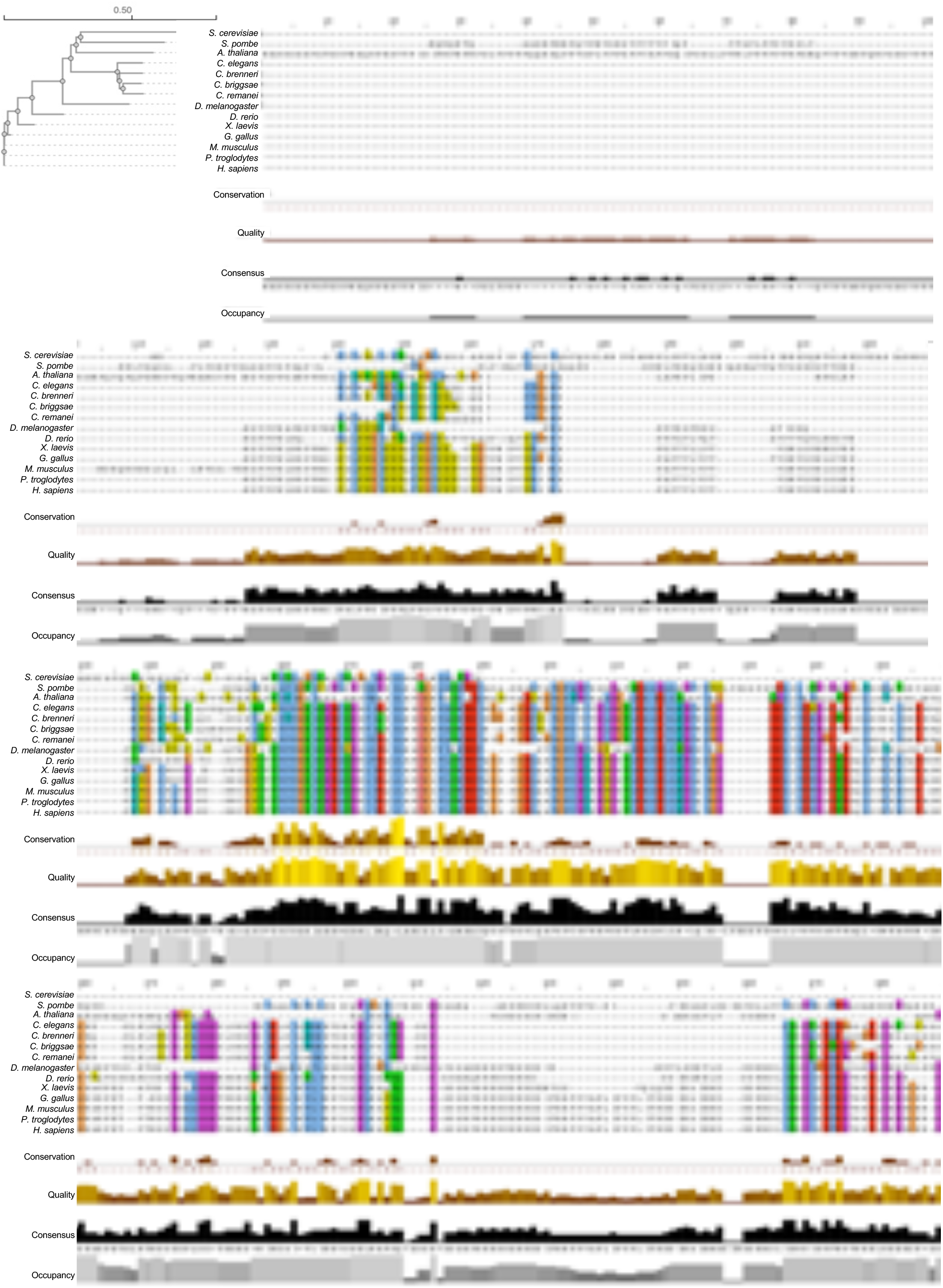
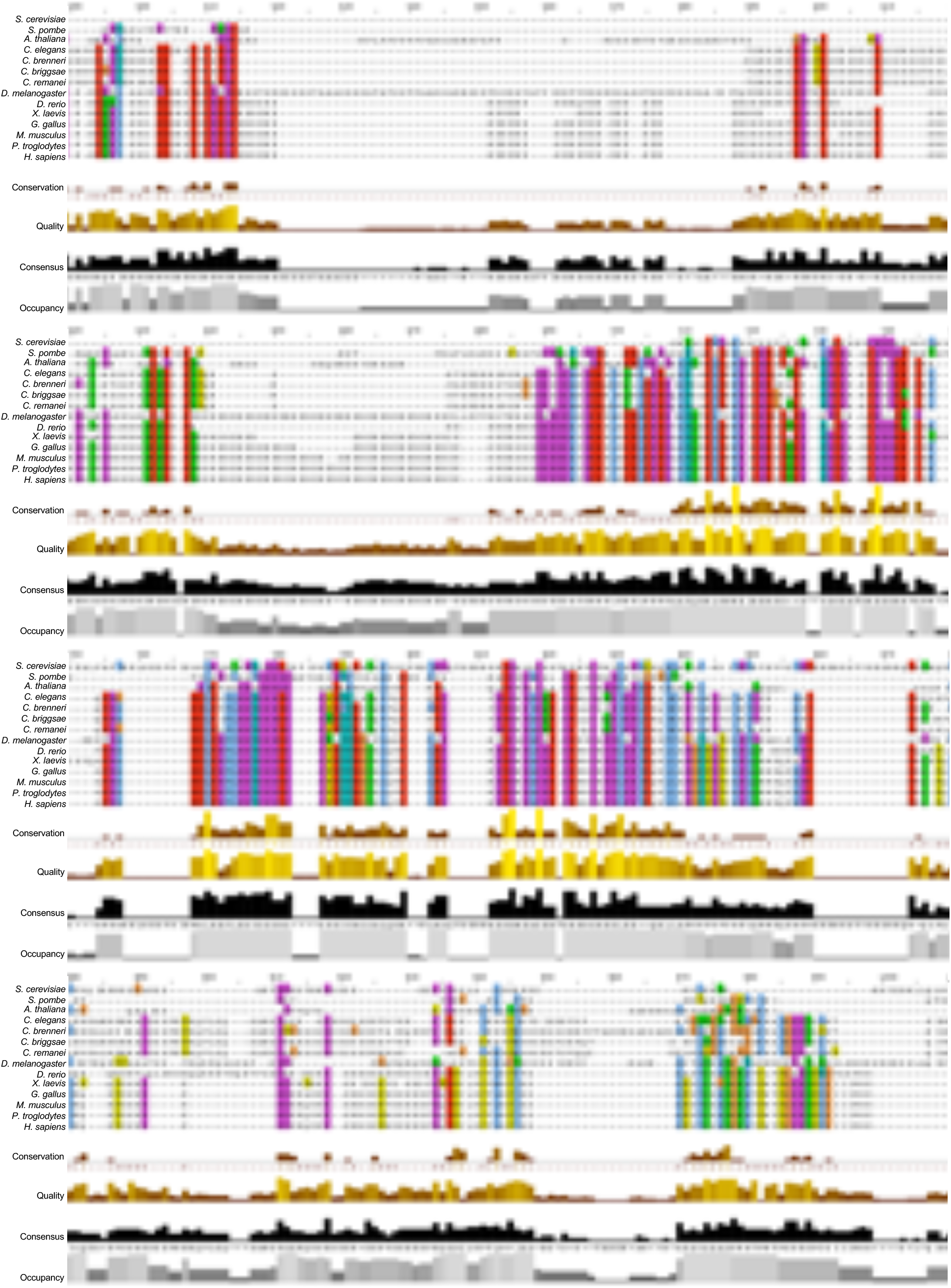
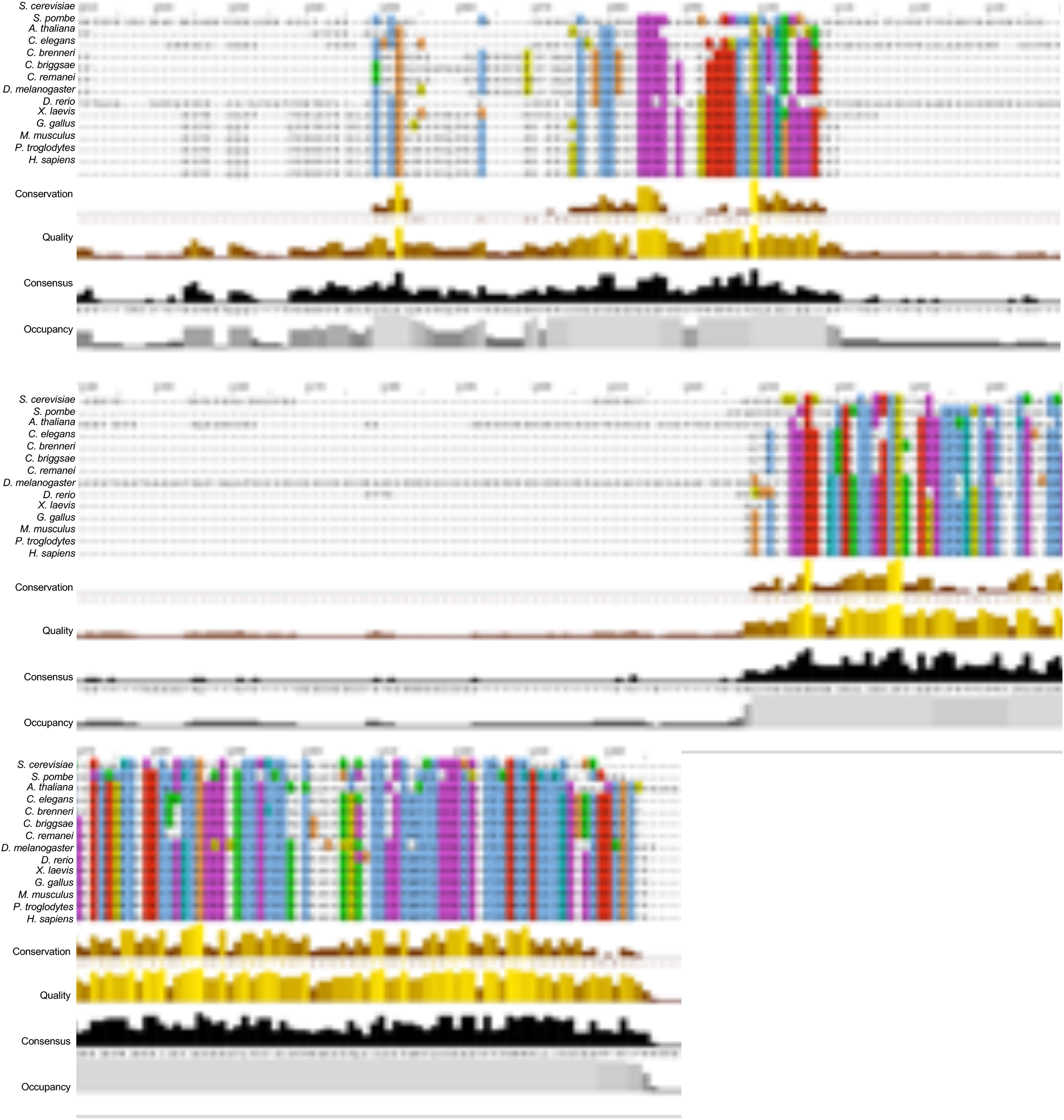
Multiple sequence alignment and phylogenetic analysis of RBM-25 orthologs. Orthologous RBM-25 protein sequences across 14 eukaryotic model organisms were retrieved from OrthoDB (Zdobnov et al., 2021), aligned using Clustal Omega, and visualized in Jalview (Waterhouse et al., 2009). The embedded phylogenetic tree was constructed using Clustal Omega (Sievers et al., 2011); the scale bar represents 0.05 amino acid substitutions per site, where a branch of this length corresponds to 5% sequence divergence between sequences.

**Supplemental Figure S7:**
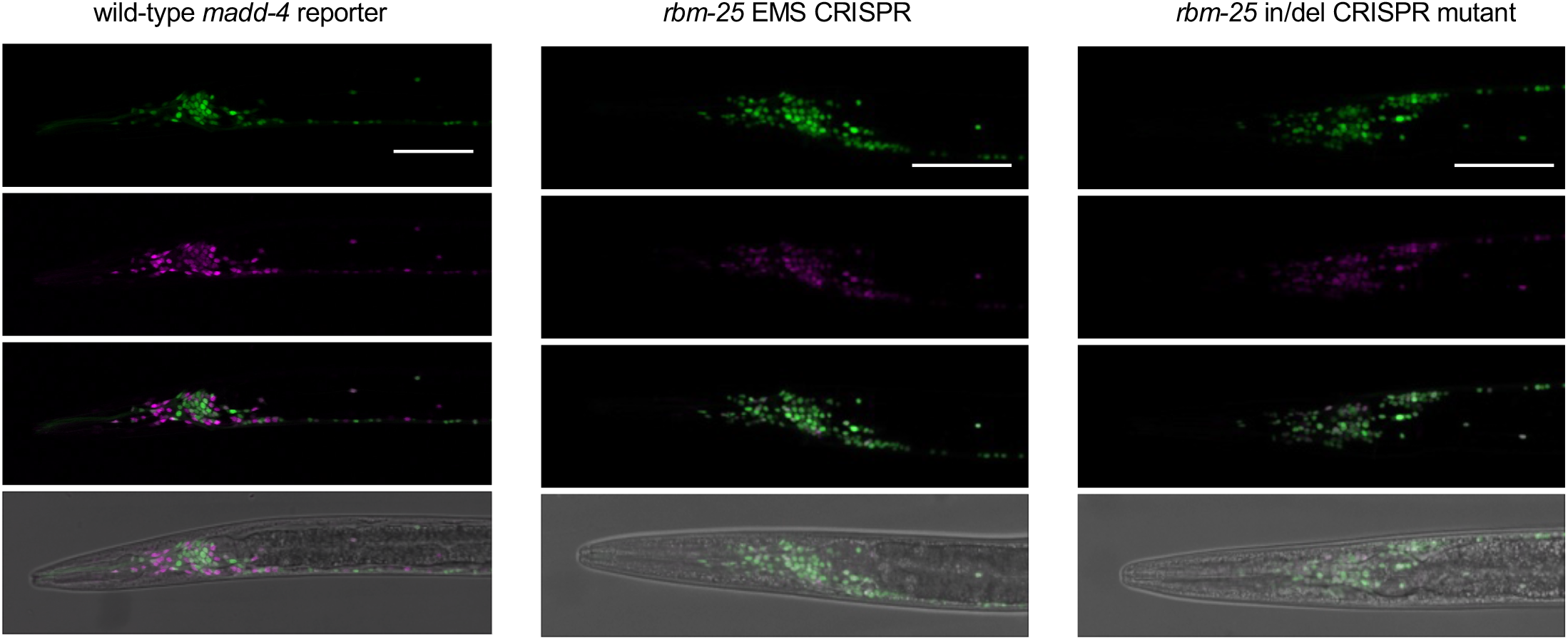
Additional alleles of *rbm-25* lead to microexon splicing defects. The *rbm-25* EMS mutant carried a point mutation that was subsequently recreated using CRISPR–Cas9 genome editing (*rbm-25* EMS CRISPR, middle panels) to assess whether this single-nucleotide change was sufficient to disrupt microexon inclusion. Additionally, we generated a *rbm-25* loss-of-function allele targeting the start codon, which introduced a frameshift and premature termination codon (PTC) at amino acid 49 (*rbm-25* in/del CRISPR, right panels). Each of these alleles recapitulated *madd-4* microexon two-colour reporter splicing defects.

**Supplemental Figure S8:**
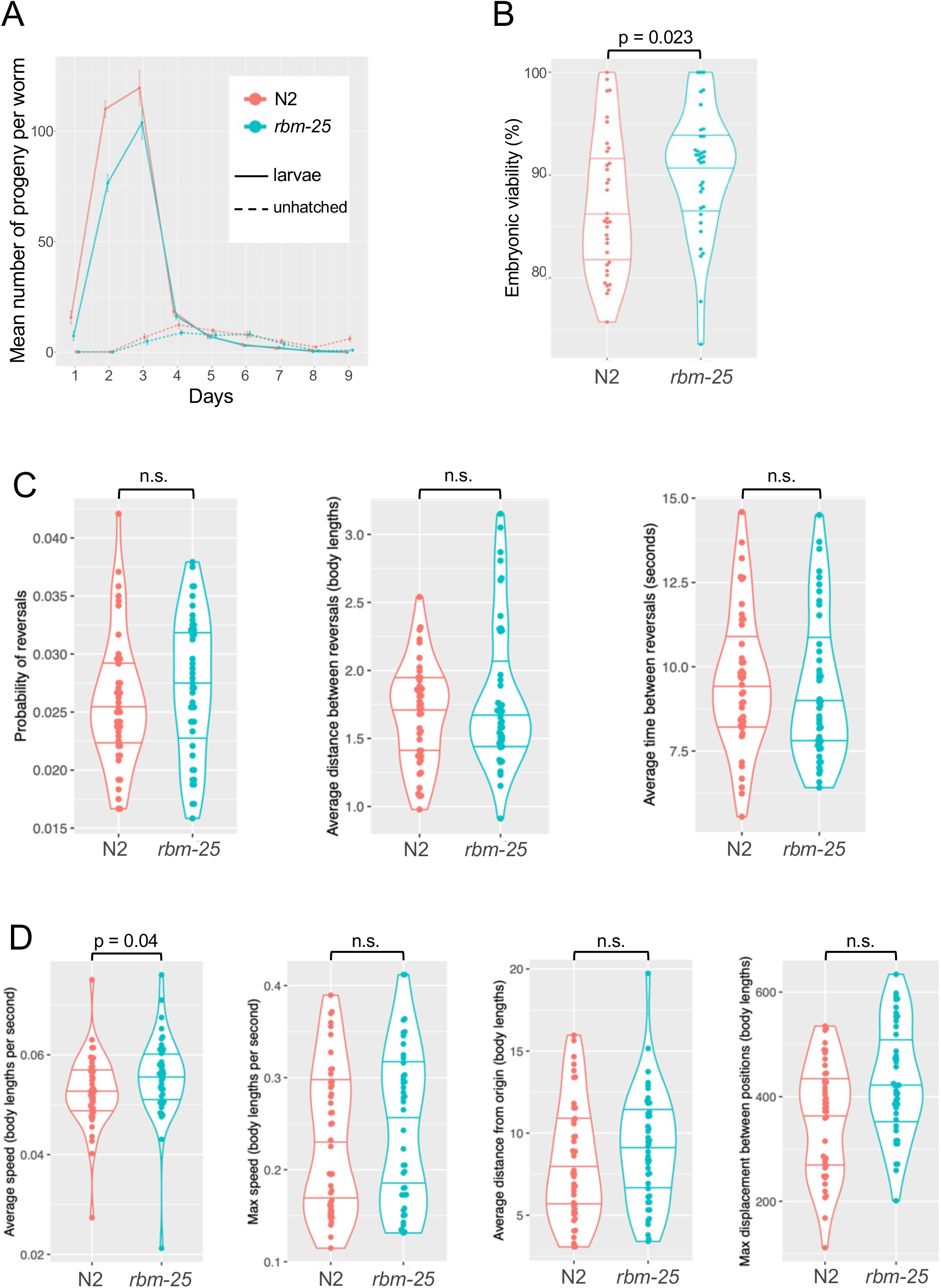
Fitness, lifespan, and locomotory responses in r*bm-25* mutants. A) Brood assay for measuring the daily output of progeny per worm in N2 and *rbm-25* worms. Mean cumulative brood size per worm was 324 (SD = 59.8) for N2 and 241 (SD = 43.1) for rbm-25. Assays were performed in duplicate with 16–20 animals per strain, for a total of n = 35 per strain after outlier removal. B) Embryonic viability measured as the percentage of laid eggs hatching into viable larvae in N2 and *rbm-25* worms. Mean viability was 87.0% (SD = 6.75%) for N2 and 90.4% (SD = 6.12%) for *rbm-25*. Differences between genotypes assessed using Wilcoxon rank-sum test. p = 0.023. n = 35 per strain after outlier removal. C) Reversal motor behaviour during locomotion on solid agar for N2 and *rbm-25* worms. Differences between genotypes assessed using Wilcoxon rank-sum test. n = 44 per strain. Probability of executing a reversal was 2.59% (SD = 0.56%) for N2 and 2.72% (SD = 0.60%) for *rbm-25*. Mean normalized distance between reversals was 1.68 body lengths (SD = 0.37) for N2 and 1.82 body lengths (SD = 0.53) for rbm-25. Mean time between reversals was 9.61 seconds (SD = 2.05) for N2 and 9.33 seconds (SD = 2.16) for *rbm-25*. D) Basic motor ability and dispersal behavior on solid agar for N2 and *rbm-25* worms. Differences between genotypes assessed using Wilcoxon rank-sum test. n = 44 per strain. Mean speed was 0.053 body lengths/s (SD = 0.007) for N2 and 0.055 body lengths/s (SD = 0.008) for *rbm-25*. Mean maximum speed over 10 minutes was 0.234 body lengths/s (SD = 0.080) for N2 and 0.246 body lengths/s (SD = 0.084) for *rbm-25*. Mean distance from origin was 8.276 body lengths (SD = 3.633) for N2 and 8.994 body lengths (SD = 3.434) for *rbm-25*. Mean maximum displacement was 359 body lengths (SD = 107) for N2 and 431 body lengths (SD = 106) for *rbm-25*.

## Notes

### Competing Interest Statement

The authors have declared no competing interest.

### Funding

- Canadian Institutes of Health Research
- Natural Sciences and Engineering Research Council

